# tRNA fragments (tRFs) populations analysis in mutants affecting tRNAs processing and tRNA methylation

**DOI:** 10.1101/869891

**Authors:** Anahi Molla-Herman, Margarita Angelova, Clément Carré, Christophe Antoniewski, Jean-René Huynh

## Abstract

tRNA fragments (tRFs) are a class of small non-coding RNAs (sncRNAs) derived from tRNAs. tRFs are highly abundant in many cell types including stem cells and cancer cells, and are found in all domains of life. Beyond translation control, tRFs have several functions ranging from transposon silencing to cell proliferation control. However, the analysis of tRFs presents specific challenges and their biogenesis is not well understood. They are very heterogeneous and highly modified by numerous post-transcriptional modifications. Here we describe a bioinformatic pipeline to study tRFs populations and shed light onto tRNA fragments biogenesis. Indeed, we used small RNAs Illumina sequencing datasets extracted from wild type and mutant *Drosophila* ovaries affecting two different highly conserved steps of tRNA biogenesis: 5’pre-tRNA processing (RNase-P subunit Rpp30) and tRNA 2’-O-methylation (CG7009 and CG5220). Using our pipeline, we show how defects in tRNA biogenesis affect nuclear and mitochondrial tRFs populations and other small non-coding RNAs biogenesis, such as small nucleolar RNAs (snoRNAs). This tRF analysis workflow will advance the current understanding of tRFs biogenesis, which is crucial to better comprehend tRFs roles and their implication in human pathology.

## INTRODUCTION

Transfer RNAs (tRNAs) are molecules of ^~^75nt transcribed by RNA polymerase III that adopt a typical cloverleaf secondary structure. They are ancient molecules required for protein translation and are encoded by hundreds of genes (^~^300 in *Drosophila*, ^~^400 in humans) often localized in clusters throughout the genome in some species (Haeusler and Engelke, 2006; Willis and Moir, 2018). Once transcribed, tRNA precursors (pre-tRNAs, ^~^125nt) are processed by the highly conserved ribozymes RNAse P and Z, to cleave the 5’ leader and the 3’ trailer, respectively (Jarrous, 2017). Then, a CCA trinucleotide tag is added at the 3’ end of mature tRNAs by a specific enzyme (RNA polymerase ATP(CTP):tRNA nucleotidyltransferase) present in all kingdoms of life. CCA tag plays a role in tRNA amino-acylation, in tRNA export towards the cytoplasm, and it has been recently suggested to participate in tRNA quality control (Wellner et al., 2018). RNase P is formed by one RNA molecule and several protein subunits such as Rpp30, which is highly conserved through evolution (Jarrous, 2017). In some species, RNAse P can also cleave non-canonical targets such as rRNA, snoRNA, some long non-coding RNA and RNAs containing N6-methyladenosine (m^6^A) (Coughlin et al., 2008; Jarrous, 2017; Park et al., 2019). Importantly, tRNA biogenesis involves the production of small RNA molecules derived either from tRNA precursors or from cleavage of mature tRNAs, hereafter referred to as tRNA fragments (tRFs). tRFs are found in a wide variety of organisms and are associated with several pathologies such as cancer and neurodegeneration (Goodarzi et al., 2015; Balatti et al., 2017; Schorn and Martienssen, 2018; Shen et al., 2018). Despite recent efforts to describe tRFs populations (Kumar et al., 2015; Schorn et al., 2017; Kuscu et al., 2018; Liu et al., 2018; Siira et al., 2018; Zhu et al., 2018; Guan et al., 2019), we are still lacking a common and versatile procedure, easy to use by the scientific community in a wide range of model systems. Thus, published tRFs analyses remain difficult to compare.

The impact of tRFs levels in various biological processes is currently under investigation. However, multiple processes impacted by tRFs have already been identified, amongst which stand translation control, transposon silencing, RNA processing, cell proliferation and DNA damage response modulation (Goodarzi et al., 2015; Kuscu et al., 2018; Li et al., 2018; Liu et al., 2018; Schorn and Martienssen, 2018; Shen et al., 2018; Guan et al., 2019; Su et al., 2019) When RNAse P cleaves the 5’ trailer of tRNA-precursor the resulting fragment is believed to be degraded by the ribonuclease translin–TRAX complex (C3PO) (Li et al., 2012) (Fig.1A). However, when RNase Z cleaves the 3’ trailer it forms Type-II tRFs (Rossmanith, 2012). Once mature, tRNAs can be cleaved forming Type-I tRFs, called 5’tRFs or 3’CCA tRFs, by Dicer or by other endonucleases that remain to be discovered (Shen et al., 2018). Besides, intermediate tRFs (i-tRFs) can be formed when cleavage takes place around the anticodon loop (Kumar et al., 2016; Sun et al., 2018). Also, mature tRNA molecules can be cut in 2 halves (tRNA halves ^~^35nt) and play important roles in different stress conditions, such as hypoxia or temperature changes (Shen et al., 2018). Spanner-tRFs can be formed before CCA addition, spanning the CCA editing point and transcription associated tRNA fragments (taRFs) are formed when RNA pol-III does not finish transcription properly (Siira et al., 2018).

**Fig.1.**
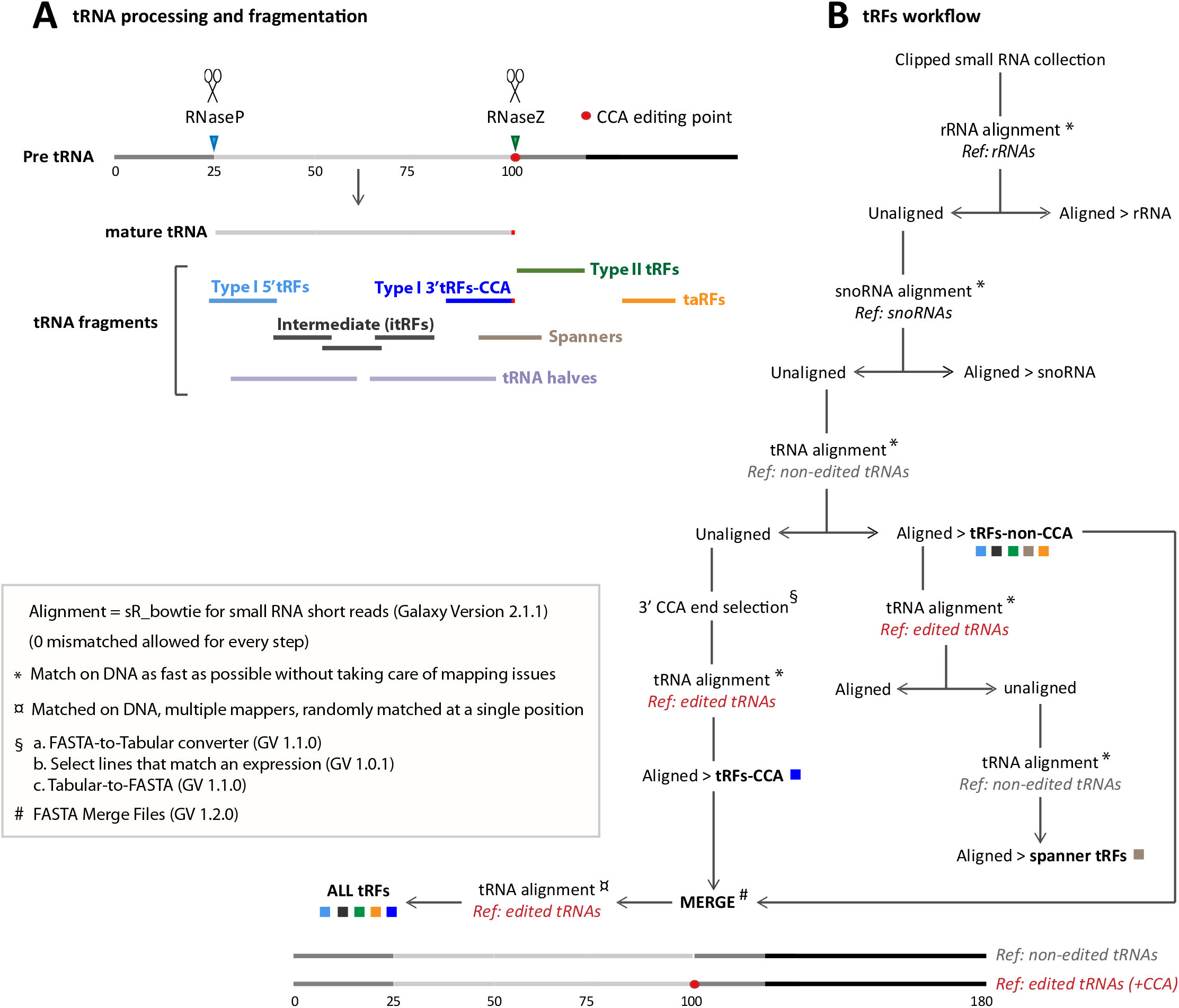
General workflow for tRNA fragments (tRFs) classes extraction: **A.** tRNA processing and tRNA fragments are depicted. The 5’ tail of Pre-tRNA is cleaved by RNase P (blue arrowhead) and 3’ tail is cleaved by RNase Z (green arrowhead). 5’ cleavage product is believed to be degraded (blue dotted lines) whereas RNase Z cleavage product forms Type-II-tRFs (green line). Mature RNA (grey) will be edited by the addition of 3’CCA motif (red). Several types of tRFs can be generated from mature RNAs, such as 5’tRFs (light blue), 3’CCA-tRFs (dark blue), and intermediate itRFs belonging to the anticodon region (black lines). Spanner-tRFs can be formed before the addition of CCA from tRNA-precursors, spanning the CCA region (light brown). Transcription associated (taRFs, orange line) can be formed from downstream regions of tRNAs. Longer tRNA halves are in light purple. **B.** Workflow developed in Galaxy to extract all classes of tRFs. All alignments were done with SR_Bowtie tool for small RNA short reads (version 2.1.1) using two types of matching: * Match on DNA as fast as possible or ¤ Match on DNA, multiple mappers. *“Ref.”* are the different genome references used for alignments in this pipeline: rRNA, snoRNA, tRNA-non-edited or tRNA-CCA-edited. To construct tRNA-non-edited reference, mature RNA (75nt) were compared with tRNA-precursors (125nt) to determine RNase P and RNase Z cleavage points. 25nt were added upstream at 5’, and 80nt downstream, right after RNase Z cleavage point (25+75+80 = 180nt approximately). To construct tRNA-CCA-edited reference we added a CCA motif to the non-edited reference, precisely at the 3’CCA edition point (red dot). tRFs CCA or non-CCA can be treated separately or altogether (ALL-tRFs).

Aberrant tRFs populations could have a *trans* effect on gene expression, similarly to other small non coding RNAs like piRNAs (Piwi interacting RNAs), which are small RNAs known to silence transposable elements (TEs) (Yamanaka and Siomi, 2015; Czech et al., 2018). Interestingly, altered tRFs have been discovered in mouse mutants for RNase Z (ELAC2) which have cardiomyopathy and premature death (Siira et al., 2018). However, it is still not known if Rnase P also plays a role in tRFs formation.

Currently, around 150-170 RNA modifications are known, and recent reports show that RNA modifications play an important role in tRFs cleavage in different organisms. Indeed, tRNA is the most extensively modified RNA in a cell (up to 25% of nucleotides per tRNA) (Delaunay and Frye, 2019; Jordan Ontiveros et al., 2019). Epitranscriptomics has recently emerged as a new field to comprehend the mechanisms underlying RNA modifications and their role in gene expression. These marks are believed to help tRNAs to respond to a wide range of environmental cues, stimuli and stress. Indeed, they play a crucial role throughout all tRNA biogenesis steps, such as sequence maturation, folding, recycling and degradation. Interestingly, there is a crosstalk between the different modification pathways and a large amount of tRNA modification enzymes defects have been linked to human pathologies (Angelova et al., 2018; Sokołowski et al., 2018; Dimitrova et al., 2019).

In *Drosophila* it has been recently shown that methylation marks protect tRNAs from cleavage and aberrant tRFs populations accumulate in mutants: on the one hand, Dnmt2 mutation impairs m^5^C methylation (Schaefer et al., 2014; Genenncher et al., 2018) and on the other hand, CG7009 and CG5220 mutation impairs 2’-O-methylation (M. Angelova, 2019). 2’-O-methylation is one of the most common RNA modifications and consists in the addition of a methyl group to the 2’ hydroxyl of the ribose moiety of a nucleoside, being also known as Nm. It is found in tRNAs, rRNAs, snRNAs (small nuclear RNAs) at the 3’ end of some small non-coding RNAs (such as piRNAs) and at some sites on mRNAs (Jordan Ontiveros et al., 2019). This modification plays a wide range of roles in RNA structure, stability and interactions (Dimitrova et al., 2019). It has been recently shown that *Drosophila* proteins CG7009 and CG5220 are the functional orthologues of yeast TRM7 (Pintard et al., 2002) and human FTSJ1 (Guy et al., 2015) respectively, which are involved in 2’-O-methylation of the anticodon loop of several conserved tRNAs substrates (tRNA-Leu, Trp, Phe) in different species, including *Drosophila*. Indeed, mutations of these tRNAs methyltransferases in *Drosophila* lead to lifespan reduction, small non-coding RNA pathways dysfunction and increased sensitivity to RNA virus infections, besides tRFs accumulation (M. Angelova, 2019).

Despite their abundance, only a very limited subset of RNA modifications can be detected and quantified by current powerful high-throughput analytical techniques such as ARM-seq, and big efforts are being invested for the development of this field (Cozen et al., 2015; Dai et al., 2017). Indeed, modifications such as 2’-O-methylation impact on classical sequencing techniques during library preparation (RT blocking), introducing a biases in the analyses (Motorin and Helm, 2019). Nevertheless, thousands of small RNA datasets have been already generated with Illumina sequencing techniques and are available for a wide range of mutants from different species (www.ebi.ac.uk/ena and www.ncbi.nlm.nih.gov/geo). Since tRFs biogenesis remains obscure, we developed and described a “user-friendly” pipeline based on Galaxy environment to study all types of tRFs, including those that have a CCA tag. To do so, we took advantage of several *Drosophila* datasets generated in our laboratories: *Rpp30* mutants, which affect tRNA processing, and *CG7009* and *CG5220*, which affect tRNA methylation (Molla-Herman et al., 2015; M. Angelova, 2019). We believe that this study will help to better understand the known pathways of tRFs biogenesis as well as to uncover new tRFs biogenesis factors and unexpected crosstalks between different RNA regulatory mechanisms, crucial for gene expression.

## MATERIALS AND METHODS

### Fly stocks

Fly stocks are described in (Molla-Herman et al., 2015; M. Angelova, 2019).

### RNA extraction from ovaries

RNA was extracted from Drosophila ovaries following standard methods detailed in (Molla-Herman et al., 2015; M. Angelova, 2019).

### Small RNA sequencing

RNA samples of 3-5 μg were used for High-throughput sequencing using Illumina HiSeq, 10% single-reads lane 1×50 bp. (Fasteris). 15–29 nt RNAs sequences excluding rRNA (riboZero) were sequenced. All the analyses were performed with Galaxy tools https://mississippi.snv.jussieu.fr. Data set deposition is described in (Molla-Herman et al., 2015; M. Angelova, 2019). European Nucleotide Archive (ENA) of the EMBL-EBI (http://www.ebi.ac.uk/ena), accession numbers: PRJEB10569 (Rpp30 mutants), PRJEB35301 and PRJEB35713 (Nm mutants).

### Clipping and concatenation

Raw data were used for clipping the adaptors [Clip adapter (Galaxy-Version 2.3.0, owner: artbio)] and FASTQ quality control was performed (FastQC Read Quality reports (Galaxy-Version 0.72)). Since replicates were homogeneous in quality and analysis (replicates for *CG7009\** heterozygous and homozygous, triplicates for *CG7009*-CG5220\** double mutants) we merged them [Concatenate multiple datasets tail-to-head (Galaxy-Version 1.4.1, owner: artbio) to have single fasta files. *CG7009*/Def9487* was used to obtain normalization numbers but is not shown in the figures.

### Data normalization using DeSeq miRNA counts

Data were normalized with bank Size Factors (SF) obtained by using [DESeq geometrical normalization (Galaxy-Version 1.0.1, owner: artbio)] with miRNA counts obtained using [miRcounts (Galaxy-Version 1.3.2)], allowing 0 mismatch (MM). Then, 1/SF values were used in Galaxy small RNA maps (Sup.Fig. 5).

### Data normalization with DeSeq using tRFs counts

To create tRFs expression heatmaps, all-tRFs read counts were normalized using [DESeq Normalization (Galaxy-Version 1.0.1, owner: artbio)] giving rise to a Normalized Hit Table.

### Genome references

rRNA, snoRNA, miRNA, ncRNA, intergenic, genic references and Transposable Elements (Ensemble canonical TE) were obtained from Ensembl Biomart (http://www.ensembl.org/biomart/martview). For tRNAs genome references, we created a genome reference of extended pre-tRNAs adding 25nt upstream and 80nt downstream of tRNA genome annotations. These sequences referred to as “non-edited tRNAs” have an average length of ^~^180.3nt (Standard Deviation 14.9nt) for nuclear tRNAs and ^~^170nt (Standard Deviation 6.2nt) for mitochondrial tRNAs. Sixteen tRNAs sequences have an intron that has to be spliced. To analyze tRFs carrying 3’CCA motif we informatically inserted a CCA in the genomic precursor sequence, at the position where tRNAs are edited after pre-tRNA maturation. We called this reference “CCA-edited-tRNAs”. We split the snoRNA sequences in two reference sets, one with box C/D snoRNAs whose mature sequence is equal or less than 120 nt long, the other with box H/ACA snoRNAs whose mature sequence is more than 120 nt long.

### small RNA annotation

Small RNA reads files were first depleted from rRNA by discarding reads aligning to rRNA genome reference. Then, we annotated the small RNAs by iterative alignments to various references using the tool [Annotate smRNA dataset (Galaxy-Version 2.4.0, owner: artbio)] and allowing 0 mismatches. Iterative alignments were performed in the following order: tRNA, tRNA-CCA-edited, miRNA, TE-derived, all-ncRNA, all genes and all intergenic. Number of alignments for each class were visualized with Pie-Charts whose sizes reflect the respective depth (total aligned reads) of the libraries (see Sup.Fig.5).

### tRFs classes extraction

Small RNA reads trimmed off from their adapter sequences were first aligned to the rRNA reference using the Galaxy tool [sR_bowtie (Galaxy-Version 2.1.1, owner: artbio)] and the option “Match on DNA as fast as possible”. Unaligned reads were retrieved and aligned to the snoRNA reference, and snoRNA alignments were visualized using the tool [small RNA maps (Galaxy-Version 2.16.1, owner: artbio)].

Next, unaligned reads were retrieved and realigned to the non-edited tRNA reference. Matching reads in this step correspond to tRFs without CCA (tRF-non-CCA) including 5’ tRFs, Type-II, spanners and intermediate tRFs. On the contrary, edited 3’CCA tRFs did not match in this step, because the CCA motif is not encoded in the genome and we did not allow mismatches (see below). To retrieve these unmatched tRFs, we selected unaligned reads with 3’ end CCA and realigned these reads to the CCA-edited-tRNA reference.

Finally, we merged non-CCA tRFs and 3’CCA-tRF using the tool [FASTA Merge Files (Galaxy-Version 1.2.0)] and realigned those reads to the CCA-edited-tRNA reference. Matched reads (“all-tRFs”) were visualized (see Fig.1) using the tool [small RNA maps (Galaxy-Version 2.16.1, owner: artbio)].

In order to isolate spanner tRFs, aligned non-CCA-tRFs were realigned using CCA-edited tRNAs as reference. Unaligned reads in this step are tRFs that span the editing point. These reads were realigned using non-edited-tRNA reference, allowing to retrieve spanner-tRFs maps.

We could not reliably detect tRNA Halves (>30nt) since our libraries were prepared using size RNA selection between 15 and 29 nt.

### tRFs global size distribution, coverage and tRF Logo

All-tRFs, non-CCA-tRFs or 3’CCA-tRFs datasets were used to generate small RNA maps and read size distributions taking into account the normalization factors for the different genotypes. Read coverage of tRNA sequences was generated using the tool [BamCoverage (Galaxy-Version 3.1.2.0.0, owner: bgruening)]. Briefly, we first used sR_bowtie with the options “matched on DNA, multiple mappers randomly matched at a single position” and “0 mismatch allowed” and tRNA-CCA-edited as a reference. Bam alignment files from this step were used with the BamCoverage tool to generate BigWig coverage files, using the library normalization factors as scale factors. The tool [computeMatrix (Galaxy-Version 3.1.2.0.0, owner: bgruening)] was then used to prepare the data for plotting heatmaps or a profile of given regions. We used four Bed files with this tool to visualize Nuclear tRNAs, 5’tRFs, 3’CCA tRFs and Mitochondrial tRNAs (see Sup.Fig.6). To obtain a Logo, tRFs FASTA files were treated to obtain last 15nt of every sequence then we used the tool [Sequence Logo (Galaxy-Version 3.5.0, owner: devteam)].

### tRFs expression heatmap and ratio calculation

To visualize tRFs expression levels we created Heatmaps. With all-tRFs collection list, we used sR_Bowtie (for small RNA short reads Galaxy-Version 2.1.1, matched on DNA, multiple mappers, randomly matched at a single position, 0 MM allowed) and we used tRNA-CCA-edited as reference. Then we used the tool [Parse items in sR_Bowtie alignment (Galaxy-Version 1.0.6)]. We did a DESeq2 normalization of hit lists (geometrical method Galaxy-Version 1.0.1, see above). We cut columns from the Normalized Hit table (Galaxy-Version 1.0.2) and we used Sort data in ascending or descending order tool (Galaxy-Version 1.0.0). This way we created a table with the tRFs counts for the different genotypes. We used Plot Heatmap with high number of rows (Galaxy-Version 1.0.0) to create the expression profile. We used Log2(value+1) and Blue-White-Red colors to reflect reads from minimal to maximal expression. To detect big changes of tRFs between genotypes, we cut columns corresponding to counts of *white*- and *Rpp30^18.2^* mutants, or *CG7009*/TbSb* heterozygous and *CG7009\** homozygous mutants. We calculated the ratio of tRFs expression between them, using Compute an expression on every row tool (Galaxy-Version 1.2.0). Data were treated with Microsoft Office Excel to better observe ratio differences by using conditional formatting tool, obtaining a three color code (Blue-White-Red from minimal to maximal value). See Sup.Fig.6.

### snoRNAs global size distribution and coverage

To represent all the reads along a flattened snoRNA molecule we analyzed the Bam Coverage, by first using sR_Bowtie for small RNA short reads (Galaxy-Version 2.1.1), matched on DNA, multiple mappers, randomly matched at a single position. 0MM were allowed, using snoRNA as reference. Then BamCoverage tool generates a coverage BigWig file from a given BAM file (Galaxy-Version 3.1.2.0.0) that we normalized using the scale factors. Afterwards, Compute Matrix prepares data for plotting a heatmap or a profile of given regions (Galaxy-Version 3.1.2.0.0). We had three Bed files to plot: snoRNAs >120nt Bed file; snoRNAs <120nt Bed file; and both together. See Sup.Fig.6.

## RESULTS

### How to study different tRFs categories

In this study we have developed a Workflow using Galaxy (https://mississippi.snv.jussieu.fr) to extract all principal classes of tRFs (Fig.1 and Methods): Type-I tRFs correspond to fragments derived from mature tRNA transcripts (5’-tRFs, 3’CCA-tRFs and intermediate i-tRFs) and Type-II, formed by RNase-Z cleavage of tRNA precursors; spanner tRFs that span CCA region created before CCA addition, and transcription associated tRFs (taRFs) due to problems in transcription termination. Thanks to this pipeline we can study them separately or altogether. Intermediate tRFs formed in or around the anticodon region are considered as non-CCA tRFs.

### tRFs description in *Drosophila* ovaries

To describe tRFs in *wild type* ovaries from young flies, we first performed a cascade of annotation of small RNA populations (Fig.2A and Sup.Fig.5) to which we “bioinformatically depleted” rRNA (see Methods). A high percentage of small RNA reads correspond to transposable elements (TEs, yellow), which correspond to piRNAs or siRNA. To distinguish tRFs carrying a 3’CCA motif from non-CCA-tRFs (5’tRFs, intermediate-tRFs, spanners, taRFs and Type-II) we used two different genome references files (see below). In *white*- ovaries there are twice as much non-CCA-tRFs than 3’CCA-tRFs (Fig. 2A: 1.15% *vs* 0.52%).

**Fig.2.**
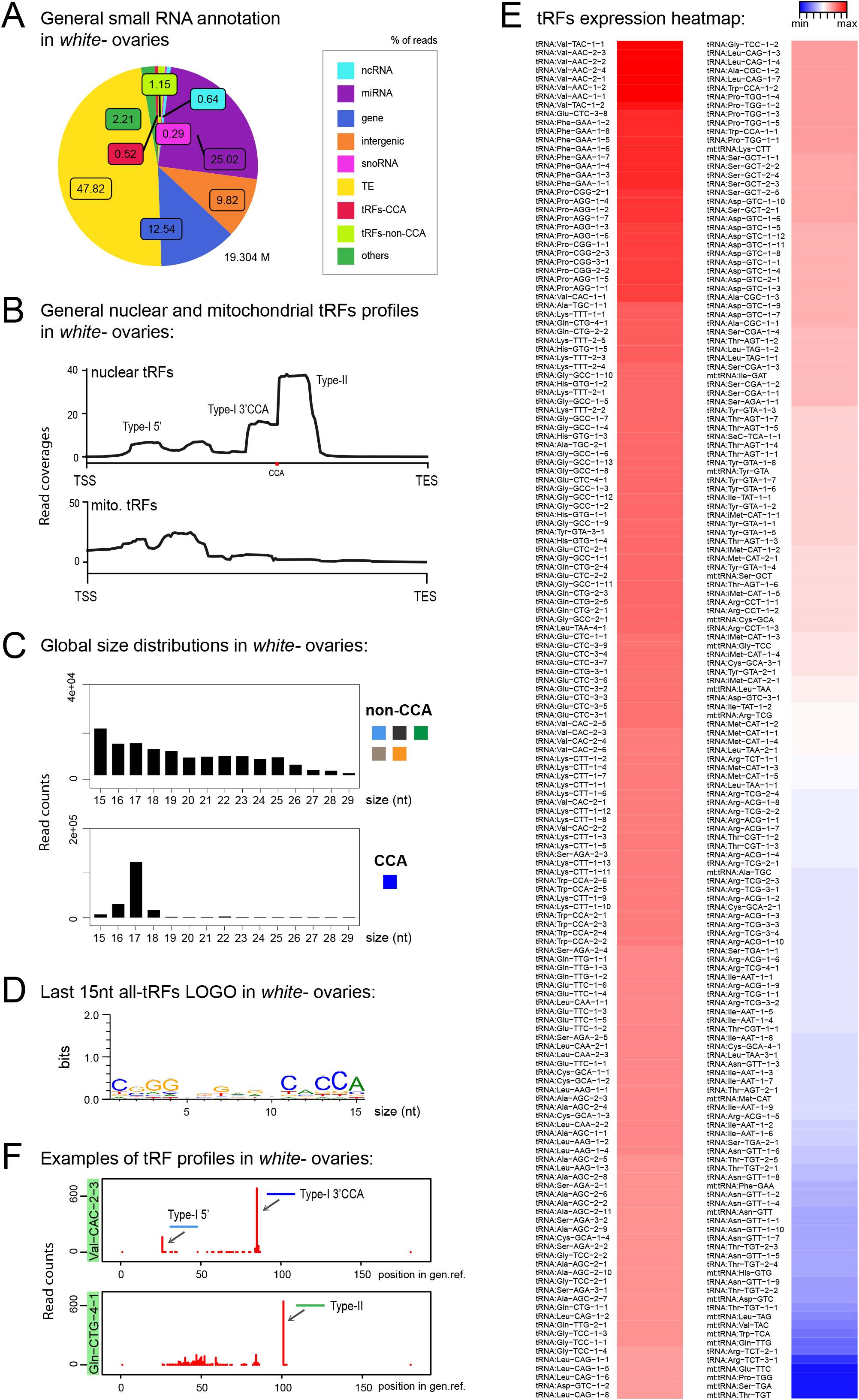
tRFs description in control *Drosophila* ovaries: **A.** Small RNAs sequences were analyzed to distinguish different categories with the help of an annotation cascade tool: miRNA, ncRNA, intergenic, genes, TE (piRNA, siRNA), snoRNAs, tRFs-non-CCA or tRFs-CCA. The percentage of reads is shown in a pie-chart, which size reflects the depth of each bank (M=Millions of reads). **B.** Nuclear and mitochondrial tRFs coverages were analyzed in *white*- control ovaries using scaling factors (see Methods). CCA edition point is shown with a red dot. Different sorts of tRFs are shown along the coverage profile from the beginning of the pre-tRNA molecule (TSS transcription start site) to the end of the extended edited genome reference (TES, transcription extended site). **C.** General size distribution (15-29nt) of normalized read counts corresponding to different categories of tRFs in *white*- control ovaries. Color-codes on the right are the same as in Fig.1B for tRFs categories. **D.** Logo for the last 15nt of *white*- tRFs sequences (all categories included, issue from *fasta* files). **E.** All-tRFs (all categories included) were counted in a hit table, normalized by DESeq normalization Geometrical method and used to do a heatmap for *white*- control ovaries, reflecting all nuclear and mitochondrial tRNA genes of *Drosophila* genome. The 3 color-code corresponds to tRFs expression, going from blue-white-red (lowest to highest levels). **F.** Examples of tRFs maps profiles in *white*- control ovaries originating from two different tRNA molecules. Red peaks reflect read counts (using scaling factors). The position of the peak along the edited tRNA genome reference reflects the beginning of the sequence. 0: beginning of the pre-tRNA. 100: position of RNase Z cleavage. Type-I 5’tRFs are in light blue, Type-I 3’CCA-tRFs are in dark blue. Type-II tRFs are in green.

Next, to have a general view of tRNA fragments, we aligned all tRFs along a canonical nuclear or mitochondrial tRNA precursor, belonging to 290 different nuclear tRNAs and 21 different mitochondrial tRNAs (Fig.2B). Nuclear tRFs coverage shows that in *white*- control ovaries there is a majority of Type-II tRFs, which corresponds to the 3’ tail processed by RNase Z. In addition, we observed a significant population of 3’CCA-tRFs and a minor population of 5’tRFs and intermediate tRFs. Mitochondrial tRFs seem more abundant at 5’ part of tRNAs molecules and around the anticodon region. In addition, global size distribution analysis showed that in control ovaries, non-CCA-tRFs are heterogeneous, ranging from 15 to 25nt, whereas 3’CCA-tRFs are mostly of 17nt long (Fig.2C). The presence of a CCA signature could be easily identified by analyzing the Logo of the last 15nt of tRFs populations (Fig.2D).

We next interrogated which type of tRNAs molecules could generate these tRFs. In *Drosophila melanogaster* there are 21 mitochondrial tRNA (one per amino-acid) and 290 nuclear tRNAs, with several tRNAs per isotype with different anticodon sequences (between 5 and 22 tRNAs per amino acid) (http://gtrnadb.ucsc.edu). For example, there are 15 tRNAs for Valine with different anticodons: 6 tRNA:Val-AAC, 7 tRNA:Val-CAC and 2 tRNA:Val-TAC. Among tRNA genes, 16 tRNAs carry an intron that needs to be spliced (tRNA:Leu-CAA, Ile-TAT and Tyr-GTA). To answer this question, we made a heatmap reflecting the expression levels of all tRFs (all types comprised) belonging to a given tRNA isotype (Fig.2E). We observed that in *white*- control ovaries, tRNA:Val-TAC or AAC were the most abundant, followed by tRNA:Glu-CTC-3-8, several tRNA:Phe-GAA and tRNA:Pro-CGG or AGG. tRNA:Val-CAC-1-1, tRNA:Ala-TGC-1-1, tRNA:Lys-TTT-1-1 or tRNA:Gln-CTG-4-1 were also abundant. Since this data comprises all tRFs categories, we described in more detail the most relevant tRFs profiles. To do so, we developed a multidimensional tRFs map which reflects the type of tRNA molecule, the read counts, and the position of tRFs along the tRNA molecule (Fig.2F). For example, in control ovaries, highly expressed tRFs from tRNA:Val-CAC-2-3 produce mostly Type-I 3’CCA-tRFs (dark blue) and Type-I 5’ tRFs, to a lesser extent (light blue). Moreover, tRNA:Gln-CTG-4-1, a tRNA which generates high amounts of tRFs in control ovaries, has a clear majority of Type-II tRFs (green).

In conclusion, our analysis describes in detail the population of tRFs present in control *Drosophila* ovaries in a global manner (annotation, coverage, size distribution, logo and heatmap tools) as well as the specific tRFs profiles of each tRNA isotype (multidimensional tRFs maps). We found that Type-II-tRFs are highly present, followed by Type-I 3’CCA-tRFs and 5’tRFs.

### tRNA processing defects lead to tRFs accumulation

We recently discovered that *Drosophila Rpp30* mutations lead to tRNA processing and early oogenesis arrest, leading to atrophied small ovaries full of early arrested stages (Molla-Herman et al., 2015). As control, we chose *white*- young (freshly hatched) ovaries described above, since they are full of early stages. Besides, we observed that *Rpp30* mutants have a defect in piRNA production. In accordance, cascade annotations showed that *Rpp30^18.2^* homozygous ovaries have decreased TE-matching sequences compared to *white*- (Fig.3A), which is rescued in *Rpp30^18.2^; ubiRpp30GFP* ovaries, showing the specificity of the phenotype. Intriguingly, we observed a substantial increase of small RNAs derived from snoRNA (pink, 6.42% in *Rpp30^18.2^* homozygous compared to 0.29% in *white*-). Moreover, we found that in *Rpp30^18.2^* homozygous ovaries, both non-CCA and CCA-tRFs were present in equal quantities (1.99 *vs* 2.09%), whereas in control ovaries non-CCA-tRFs were more expressed than CCA-tRFs (1.15% *vs* 0.52%). This suggests is an increase of CCA-tRFs and/or a decrease of some non-CCA-tRFs in *Rpp30^18.2^* homozygous mutants.

**Fig.3.**
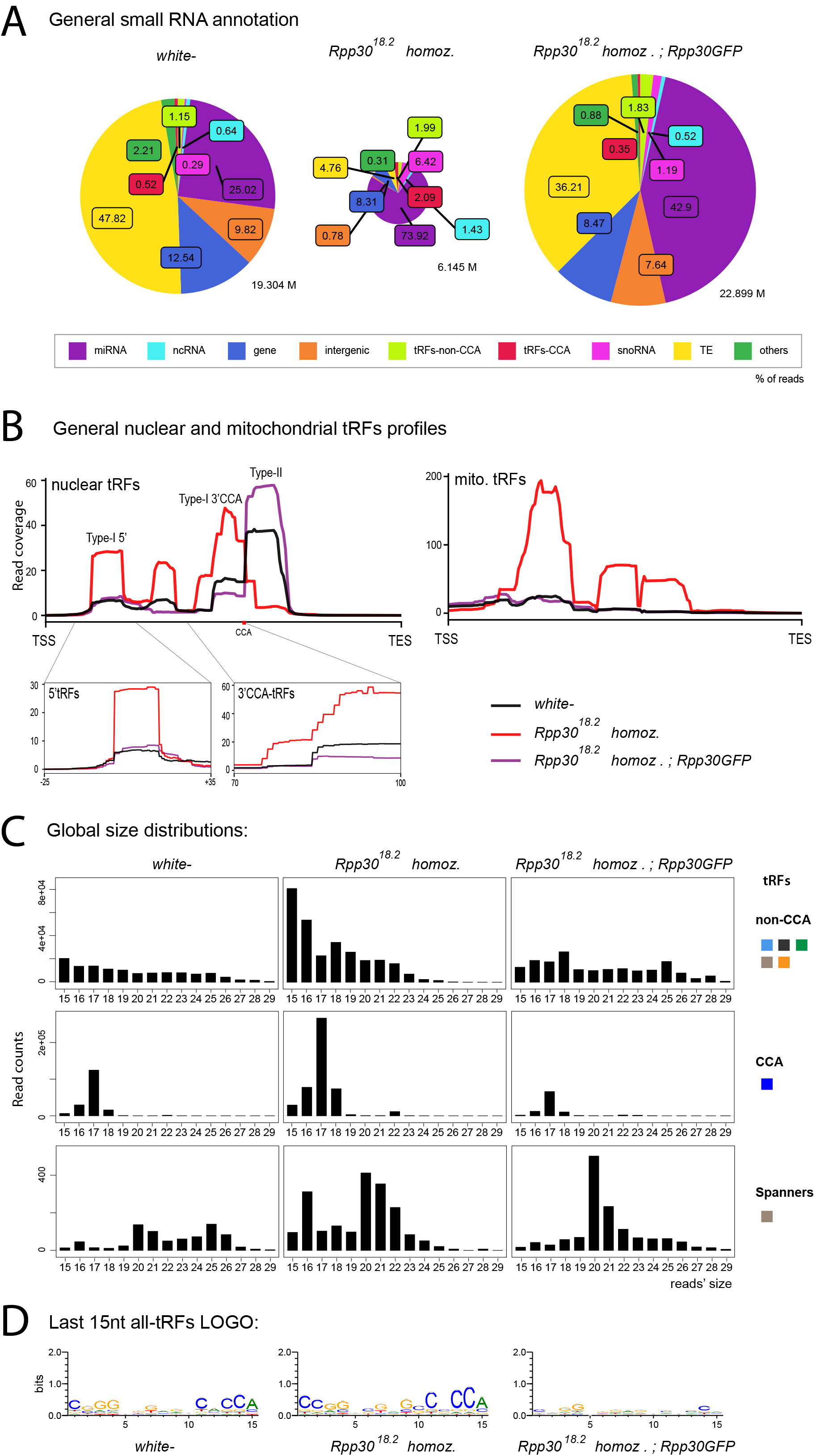
tRNA processing plays a role in nuclear and mitochondrial tRFs formation: **A.** Small RNAs sequences were analyzed in different genotypes to distinguish categories with the help of an annotation cascade tool: miRNA, ncRNA, intergenic, genes, TE (piRNA, siRNA), snoRNAs, tRFs-non-CCA or tRFs-CCA. The percentage of reads for each genotype is reflected in pie-charts, which size reflects the depth of each bank (M=Millions of reads). **B.** Nuclear and mitochondrial tRFs coverage was analyzed in *white*- control and *Rpp30* mutant ovaries using scaling factors (see Methods). Different tRFs are shown along the coverage profile from the beginning of the pre-tRNA molecule (TSS transcription start site) to the end of the extended edited genome reference (TES, transcription extended site). CCA edition point is shown in red. Type-I 5’tRFs and 3’CCA-tRFs region are zoomed in for better comparison between genotypes. **C.** General size distribution (15-29nt) of normalized read counts corresponding to different categories of tRFs in *white*- control and mutant ovaries. Color-codes on the right are the same as in Fig.1B for tRFs categories. **D.** Logo for the last 15nt tRFs sequences of *white*- control and mutant ovaries (all categories included, issued from *fasta* files containing all different sequences).

Nuclear tRFs coverage (Fig.3B, left panel) showed that in *Rpp30^18.2^* homozygous ovaries, there is a substantial increase of Type-I tRFs (5’tRFs, intermediate tRFs and 3’CCA), and a drastic decrease of Type-II tRFs when compared to control (red versus black lines). Importantly, rescued *Rpp30^18.2^; Rpp30GFP* ovaries (purple line) showed a similar profile to *white*-, demonstrating that Rpp30 overexpression is able to recover tRFs formation in *Rpp30^18.2^* homozygous mutants. In parallel, mitochondrial-tRFs coverages (Fig.3B, right) showed that *Rpp30^18.2^* homozygous have a high accumulation of different tRFs types.

Next, global size distribution (Fig.3C) indicated that tRFs accumulate in *Rpp30^18.2^* homozygous compared to *white*-. Indeed, non-CCA-tRFs range from 15 to 22nt whereas 3’CCA-tRFs are mostly of 17nt long in mutants (Fig. 3C, D). Finally, spanner-tRFs, which is a very minor population in *Drosophila white*- ovaries, are heterogeneous in size and do not show important changes in mutants when compared to control (Fig.3C, lower panels).

In conclusion, our analysis shows that tRNA processing defects alter tRFs biogenesis in *Rpp30* mutants: Type-I tRFs increase (5’tRFs, intermediate tRFs and 3’CCA tRFs), and Type-II tRFs decrease. Since there are more than 300 tRNAs genes in *Drosophila*, we wondered if these defects were due to a specific tRNA.

### tRFs expression levels are altered in *Rpp30* mutants

As mentioned, tRFs heat maps showed that *white*- control ovaries have extremely abundant tRFs derived from tRNA-Val, Glu, Phe, Pro, Ala, Lys, Gln (Fig.2E and 4A). Importantly, the general profile changes in *Rpp30^18.2^* homozygous and is partially rescued in *Rpp30^18.2^; ubiRpp30GFP* (Fig.4A). To easily detect the most drastic changes in tRFs populations we calculated the ratio of tRF-counts between *Rpp30^18^*^.2^ homozygous and *white*- ovaries (Fig.4B). For example, tRFs derived from tRNA:Val-AAC-2-2 are highly decreased in *Rpp30^18.2^* homozygous ovaries compared to *white*-, with a ratio of 0.04 (Fig.4B, blue).

**Fig.4.**
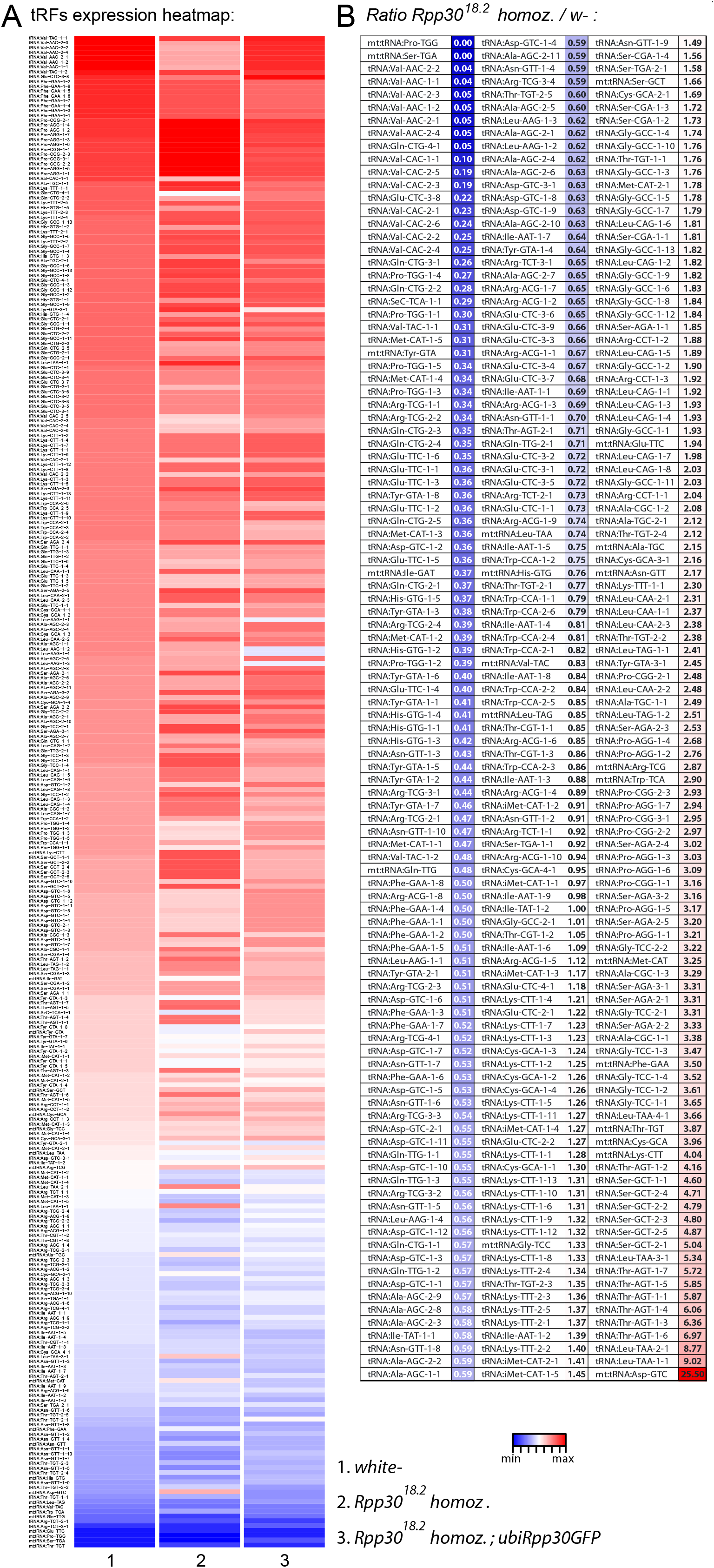
tRFs expression is altered in Rpp30 mutants: **A.** all-tRFs (all categories included) were counted in a hit table, normalized by DESeq normalization Geometrical method and used to generate a heatmap for different genotypes (1-3) reflecting all nuclear and mitochondrial tRNA genes in *Drosophila’s* genome. Counts were sorted from maximum to minimum values for the *white*- column (left) for better comparison with other genotypes. Levels of expression are reflected with a color-code going from blue (lowest levels) to white (middle levels) to red (highest levels). **B.** The ratio *Rpp30^18.2^* homozygous/*white*- of counts found in the heat table was calculated. Minimal values are reflected in dark blue, middle values in white, and maximal values in red.

From this ratio data, we selected tRNA profiles in which tRFs were increased, decreased or unchanged in mutants when compared to *white*- (Fig.5). For example, in *Rpp30^18.2^* mutants: tRNA:Leu-TAA-1-1, tRNA:Thr-AGT-1-6, tRNA:Ser-GCT-2-1, tRNA:Gly-TCC-1-2 and tRNA:Pro-AGG-1-5 show an increase of 3’CCA-tRFs. In addition, tRNA:Ala-CGC-1-1 accumulates 3’CCA-tRFs and 5’tRFs. tRNA:Ser-AGA-2-2 shows a drastic increase in only 5’tRFs. Indeed, all tRNA:Ser-AGA/CGA (12 different tRNAs) behave similarly. tRNA:Leu-CAA-2-2 has an important increase in 5’tRFs as all tRNA:Leu-CAA. It should be noted that Leu-CAA group have an intron of 40-44nt, that is why 3’CCA-tRFs are located offset in tRFs maps. Next, tRNA-Gly-GCC-2-1 is similar in *white*- and *Rpp30^18.2^* mutants. Finally, several tRFs types decreased in *Rpp30^18.2^* mutants: Type-II-tRFs generated from tRNA:Glu-CTC-3-8 and tRNA:Gln-CTG-4-1; 5’-tRFs generated from tRNA:Val-CAC-2-3; 3’CCA-tRFs generated from tRNA:Val-CAC-2-2 and 2-3. We also compared tRFs profiles by selecting the mostly expressed tRNAs in *white*- and we compared them to mutants (Sup. Fig. 1).

**Fig.5.**
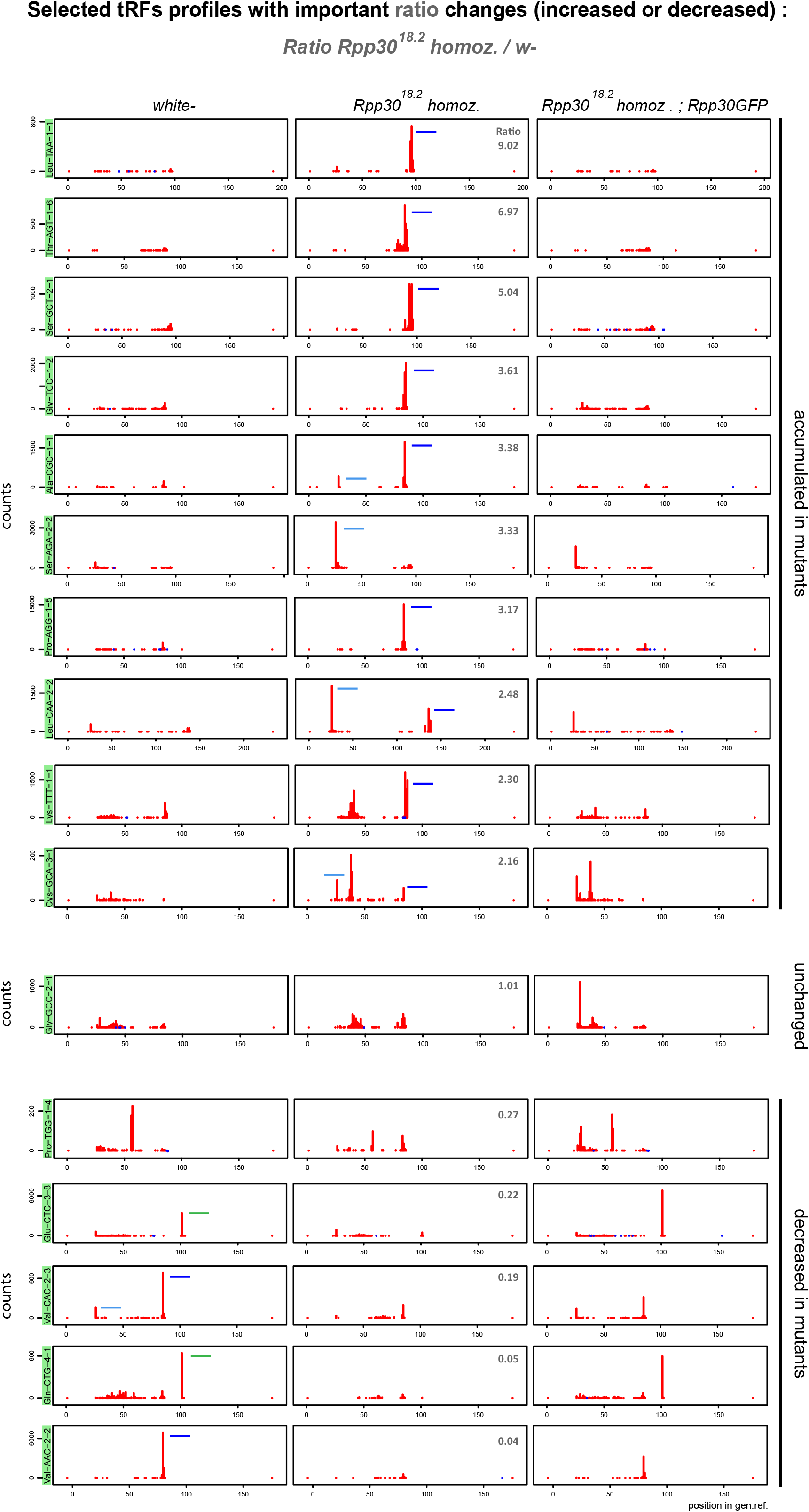
Rpp30 mutation leads to an increase of 5’tRFs and 3’CCA-tRFs but a decrease of Type-II-tRFs. 16 tRFs maps profiles belonging to examples of the most changed (increased or decreased) tRFs from *Rpp30^18.2^ homoz. /white*- ratio (see in Fig.4B) are shown for the different genotypes, using normalizing factors (see Methods). Since pre-tRNAs sequences are included in the tRNA genome reference, 5’tRFs are located at the position 25nt instead of position 0nt. 3’CCA-tRFs are located around the position 75nt and Type-II RNase Z cleavage product tRFs are located around position 100nt, depending on the length of the tRNA and if they have an intron. The peak determines the beginning of the sequence. tRFs are schematized in *white*- and *Rpp30^18.2^* homozygous for better comparison: 5’tRFs in light blue, 3’CCA in dark blue and Type-II in green. Ratio’s values above 1 : tRFs increased in *Rpp30^18.2^* mutants. Ratio’s values below 1: tRFs decreased in *Rpp30^18.2^* mutants.

Overall we find that in *Drosophila* ovaries, tRFs originate from most isotypes of tRNAs and show heterogeneous profiles that can change even within the same tRNAs isotype in mutant conditions. In general, as shown in Fig.3B, we find that Type-II-tRFs decreased in *Rpp30* mutants, whereas Type-I tRFs accumulated. tRNA processing by RNase P is the first step of tRNA biogenesis following transcription. We thus wondered whether other downstream events could also affect tRFs biogenesis, such as tRNA post-transcriptional modifications of tRNA molecules.

### tRNA methylation defects lead to a decrease of Type-II tRFs and an increase of 3’CCA-tRFs

Mutations of tRNAs 2’-O-methyltransferases (Nm MTases) CG7009 and CG5220 lead to *Drosophila* life span reduction, small RNA pathways dysfunction, increased sensitivity to RNA virus infections and tRFs-Phe accumulation (M. Angelova, 2019). In our new analysis, non-CCA-tRFs decrease in *CG7009\** homozygous mutants when compared to control (Fig.6A, light green), whereas tRFs-CCA show no significant difference in the three tested genotypes (Fig.6A, red). Surprisingly, double mutants *CG7009*, CG5220\** show profiles similar to control. The analysis of tRFs size distribution using normalization factors and of a Logo sequence revealed that 3’CCA-tRFs of 18nt increase in *CG7009\** homozygous mutants when compared to control (Fig.6B and C), which was rescued in double mutants. This rescue suggests that CG5220 Nm methylation at position 32 of tRNA might be important for the stabilization of the increased 3’CCA-tRFs observed in *CG7009\** homozygous ovaries. Finally, lowly expressed spanner tRFs were similar in *CG7009\** heterozygous and homozygous mutants when compared to control and slightly lower in double mutants (Fig.6B).

**Fig.6.**
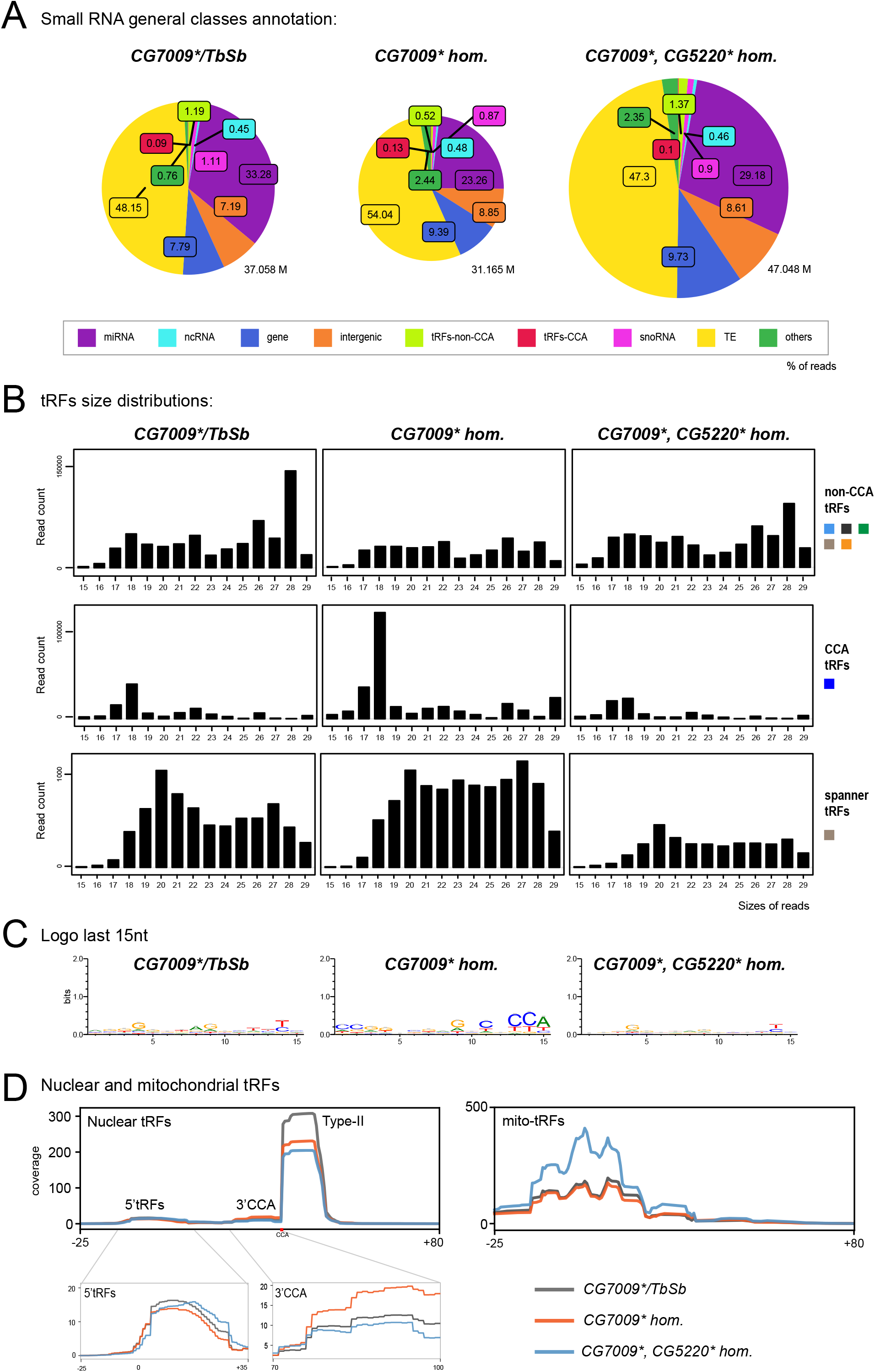
tRNA methylation defects alters nuclear and mitochondrial tRFs formation: **A.** Small RNAs sequences were analyzed in different genotypes to distinguish different categories with the help of an annotation cascade tool: miRNA, ncRNA, intergenic, genes, TE (piRNA, siRNA), snoRNAs, tRFs-non-CCA or tRFs-CCA. The percentage of reads *CG7009\** heterozygous, *CG7009\** homozygous and *CG7009*, CG5220\** double mutant ovaries is reflected in pie-charts. The pie-charts size reflects the depth of the bank (M=Millions of reads). **B.** General size distribution (15-29nt) of normalized read counts corresponding to different categories of tRFs in different genotypes using scaling factors (see Methods). Color-codes on the right are the same as in Fig.1B for tRFs categories. **C.** Logo for the last 15nt tRFs sequences of control and mutant ovaries (all categories included, issued from *fasta* files). **D.** Nuclear and mitochondrial tRFs coverages were analyzed in different genotypes using scaling factors (see Methods). Different sorts of tRFs are shown along the coverage profile from the beginning of pre-tRNA molecule (TSS transcription start site) to the end of the extended edited genome reference (TES, transcription extended site). CCA edition point is shown with a red dot. Type-I 5’tRFs and 3’CCA-tRFs region are zoomed for better comparison between the genotypes.

To obtain an overview of which tRFs classes were globally altered in Nm MTases mutants, we aligned all-tRFs together along a canonical nuclear or mitochondrial tRNA precursor. In heterozygous control ovaries (Fig.6D, grey), there is a majority of Type-II nuclear tRFs, similarly to control *white*- ovaries (Fig.2B). Of note, the size of heterozygous ovaries is bigger than *white*- ovaries, since they have early and older stages. Interestingly, *CG7009\** homozygous mutants and double mutants *CG7009*, CG5220\** showed a decrease of Type-II tRFs when compared to heterozygous control (Fig.6D, orange and blue), suggesting that these Nm MTases are involved in tRFs biogenesis.

Since Type-II-tRFs reads signal is very high and the signal of 5’tRFs and 3’CCA-tRFs is lower, it was difficult to identify major changes in Type-I tRFs. By zooming into these regions, we observed that 5’tRFs slightly decrease in *CG7009* homozygous* mutants (Fig.6D, left panel, orange), whereas 3’CCA-tRFs increase (Fig.6D, right panel, orange). Surprisingly, double mutants show similar profiles to control, suggesting that CG5220 mutation somehow rescues CG7009 defects on 3’CCA-tRFs accumulation. Interestingly, we recently reported that longer 5’tRF-Phe (^~^35nt) was significantly increased in different combinations of *CG7009* mutant alleles (M. Angelova, 2019).

Moreover, we observed mitochondrial tRFs in heterozygous ovaries similar to *white*- flies (Fig.2B and Fig.6D, right panel, grey line), derived mostly from the first half of the molecule. Homozygous mutant for *CG7009\** ovaries are similar to heterozygous mutants, whereas double mutants *CG7009*, CG5220\** show a global increase of mito-tRFs, suggesting that CG5220, and not CG7009, may be somehow involved in mitochondrial-tRFs biogenesis.

In summary, we have observed that defects of tRNA 2’-O-methylation affect tRFs populations. *CG7009* and *CG5220* mutations lead to a decrease of Type-II tRFs and *CG7009* mutation leads to an accumulation of 3’CCA-tRFs and a slight decrease of 5’tRFs.

### tRNA methylation mutations affect tRFs derived from different isotypes of tRNAs

tRFs expression heat maps showed that the most expressed tRFs in heterozygous mutant for *CG7009\** ovaries were those corresponding to tRNAs Glu-CTC, Pro-CGG and AGG, Val-TAC, Cys-GCA, Lys-TTT, Gly-TCC, Ala-CGC, His-GTG, Ser-GCT (Fig.7A), similarly to *white*- control ovaries (Fig.2E). In *CG7009\** homozygous mutants, tRFs derived from Glu-CTC, Cys-GCA, Lys-TTT or Gly-TCC decreased, whereas tRFs derived from Ser-GCT increased compared to control. These changes have been quantified as in Fig. 4B, by calculating the ratio between homozygous and heterozygous *CG7009\** reads counts.

**Fig.7.**
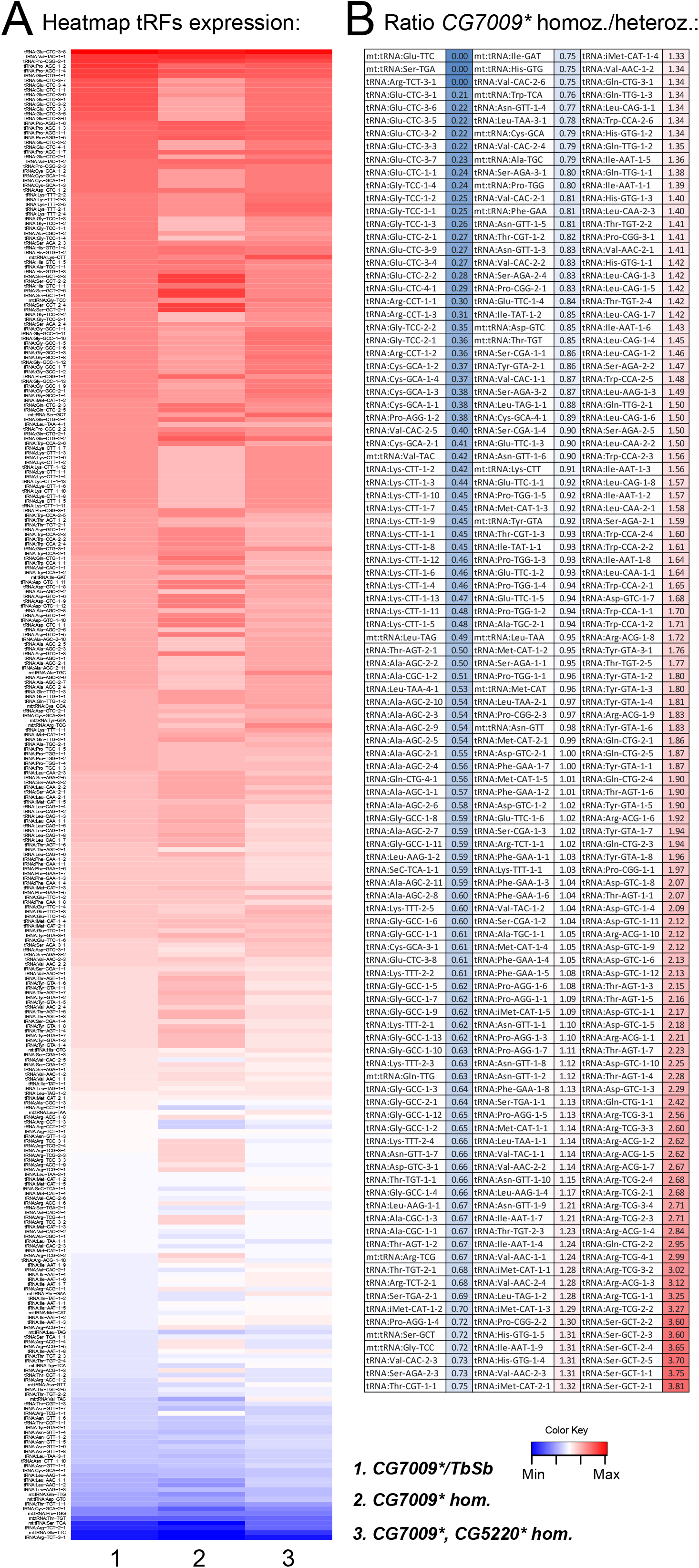
tRFs expression is altered in tRNA methylation mutants: **A.** ALL-tRFs were counted in a hit table, normalized by DESeq normalization Geometrical method and used to generate a heatmap for the different genotypes (1-3) reflecting all nuclear and mitochondrial tRNA genes in *Drosophila’s* genome. Counts were sorted from maximum to minimum values for the *CG7009*/Tb,Sb* heterozygous column (left) for better comparison with other genotypes. Levels of expression are reflected with a color-code going from blue (lowest levels) to white (middle levels) to red (highest levels). **B.** The ratio *CG7009\** homozygous/heterozygous of counts found in the heat table was calculated. Minimal values are in dark blue, middle values in white, and maximal values in red.

Considering the ratio change between homozygous and heterozygous *CG7009\** ovaries (Fig. 8, upper panels) we can observe that 5’tRFs are strongly affected for several tRNAs, such as Glu-CTC, Gly-TCC, Cys-GCA a defect that is slightly rescued in double mutants. In addition, we observe that Type-II tRFs from tRNAs Gln-CTG-4-1 and Pro-AGG-2-1 are highly affected. tRNA:Met-CAT-1-5 tRFs do not change between control and mutants, where we observe a population of tRFs that match the anticodon region. On the contrary, we clearly see an increase of 3’CCA tRFs in *CG7009\** homozygous mutants for several tRNAs: Pro-CGG-1-1, Thr-AGT-1-4, Gln-CTG-1-1, Arg-TCG-2-1, Ser-GCT-2-2. Surprisingly, those defects are rescued in double mutants *CG7009*, CG5220\**. In addition, we found similar results analyzing profiles corresponding to highly expressed tRFs in heterozygous *CG7009\** ovaries for 5’tRFs and Type-II tRFs (Sup.Fig.2). However, increase of 3’CCA-tRFs were more difficult to observe, indicating that 3’CCA tRFs increased in *CG7009\** homozygous ovaries are not highly present in heterozygous ovaries.

**Fig.8.**
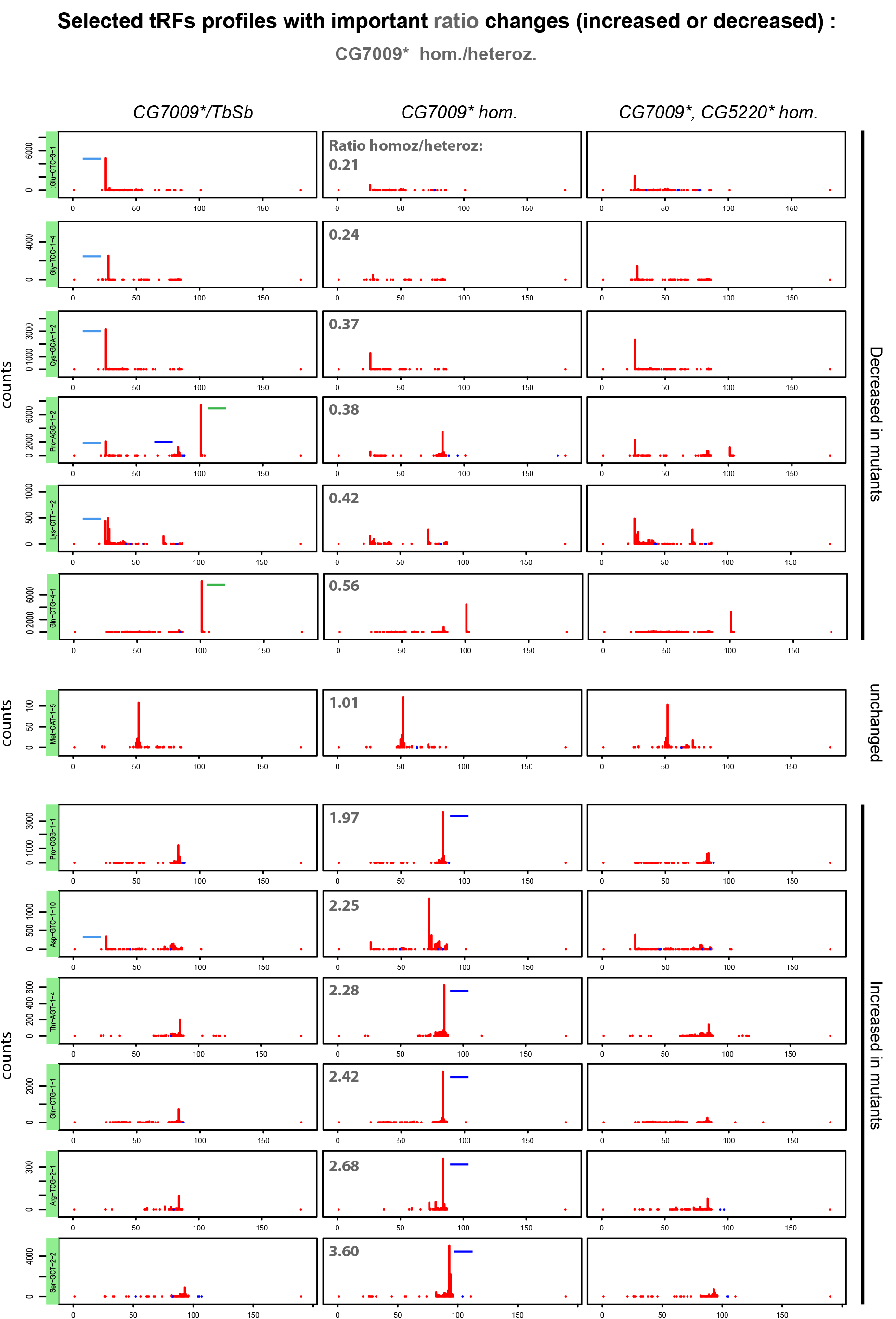
tRFs biogenesis is altered in tRNA methylation mutants: 13 tRNAs tRFs maps profiles belonging to examples of the most changed (increased or decreased) tRFs from *CG7009\** homozygous/heterozygous ratio (see Fig.7B) are shown for the different genotypes, using normalizing factors (see Methods). Since pre-tRNAs sequences are included in the tRNA-reference, 5’tRFs are located at the position 25nt instead of position 0nt. 3’CCA-tRFs are located around the position 75nt and Type-II RNase Z cleavage product tRFs are located around position 100nt, depending on the length of the tRNA and if they have an intron. The peak determines the beginning of the sequence. tRFs are schematized in *CG7009\** homozygous and heterozygous mutants for better comparison: 5’tRFs in light blue, 3’CCA in dark blue and Type-II in green. Ratio’s values above 1: tRFs increasing in *CG7009\** homozygous mutants. Ratio’s values below 1: tRFs decreasing in *CG7009\** homozygous mutants.

CG7009 and CG5220 have been shown to methylate tRNA-Leu, Trp, Phe (conserved targets in yeast and humans) and CG5220 methylates tRNA-Glu and -Gln in *Drosophila* [bioRxiv ref]. Indeed, tRFs derived of these specific tRNAs showed different profiles. (Sup.Fig.3). First, tRNA-Leu tRFs had different profiles regarding their anticodon sequence. Some showed 5’tRFs in control ovaries that are decreased in *CG7009\** mutants, and that remain similar or decrease in double mutants. Type-II tRFs decrease in *CG7009\** homozygous mutants whereas 3’CCA increase. Thus, tRNA-Leu tRFs follow the general tendency observed in Fig.6D. In contrast, tRNA-Trp tRFs show an increase of 3’CCA tRFs, however, double mutants lose 3’CCA tRFs. Finally, tRNA-Phe derived 3’CCA tRFs increased in *CG7009\** homozygous mutants, when double mutants *CG7009*, CG5220\** lose 3’CCA tRFs.

Overall, tRNA Nm methylation defects in the anticodon loop has a global impact on tRNA fragmentation, to a lesser extent than tRNA processing deffects (Fig.9). Indeed, Type-II tRFs show a decrease in *CG7009\** homozygous mutants and Type-I 3’CCA tRFs are increased, whereas 5’tRFs are slightly decreased. Intriguingly, double mutants *CG7009*, CG5220\** have profiles similar to control, indicating that for at least some of the observed differentially expressed tRFs increased in *CG7009\** homozygotes, CG5220-dependent Nm modification might have an effect on their biogenesis.

**Fig.9.**
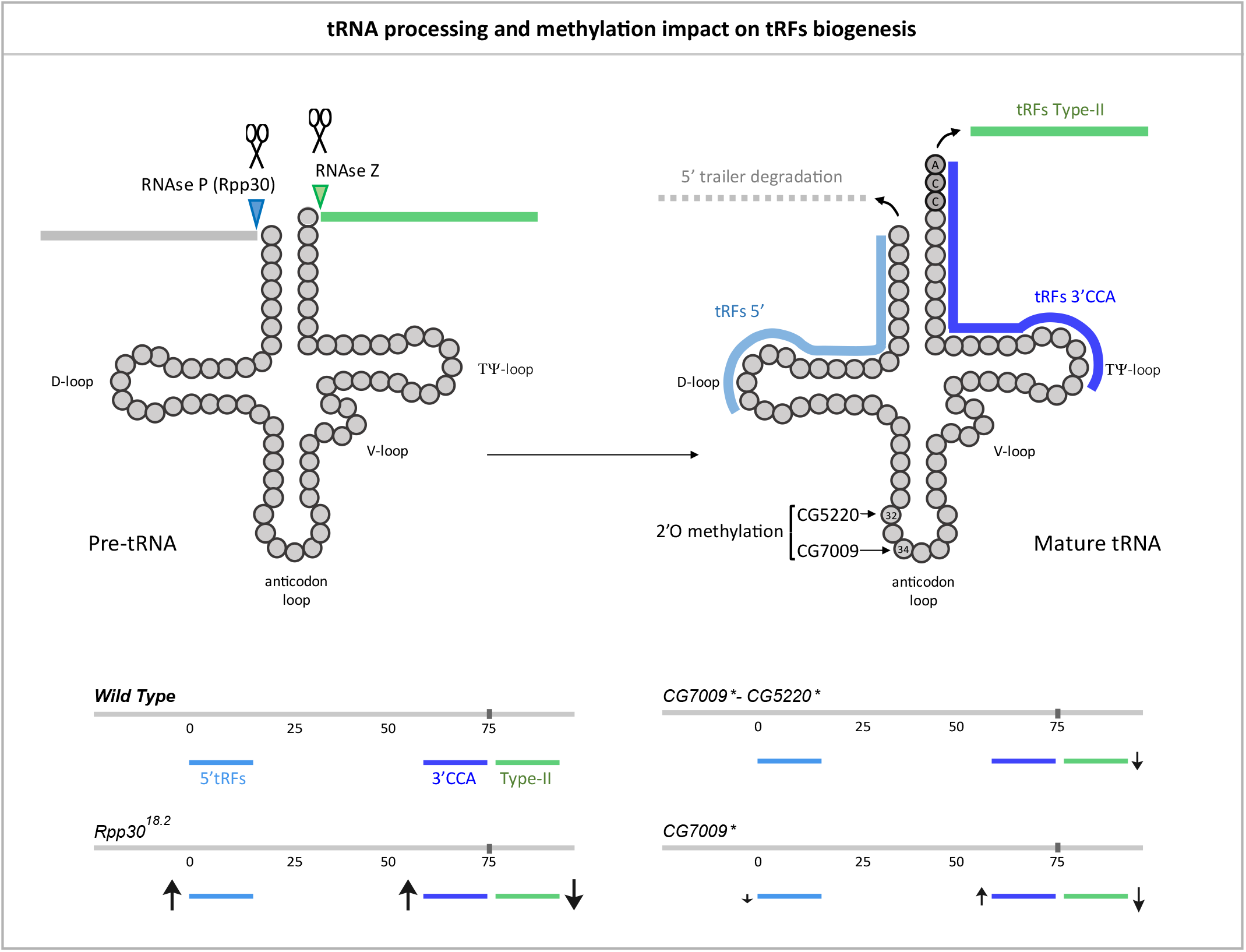
tRNA processing and methylation defects impact on tRFs biogenesis: Main steps of tRNA processing are depicted. Cleavages sites for ribozymes RNase P and Z are indicated on a pre-tRNA molecule. Cleavage of 3’ trailer forms Type-II tRFs (green). Once mature, CCA is added (dark grey) and tRNAs can be cleaved at the D-loop, forming the tRFs 5’ (light blue) and at the T-loop, forming 3’CCA tRFs. 2’O methylation sites for CG5220 and CG7009 are shown at the anticodon loop. Increase or decrease of different tRFs populations in mutants for tRNA processing or tRNA methylation are schematized with arrows of different sizes (↑, increased, ↓, decreased).

### tRNA processing and methylation avoid snoRNA fragmentation

The increase of small RNAs derived from snoRNAs observed in *Rpp30* mutants (Fig.3A) invited us to study this population in more detail. In *Drosophila*, snoRNAs >120nt are box H/ACA and play a role in Pseudouridylation whereas snoRNAs <120nt are box C/D snoRNA and play a role in 2’-O-methylation (Huang et al., 2005; Angrisani et al., 2015; Falaleeva et al., 2017). Since RNase P has been shown to participate in snoRNAs maturation in some species (Coughlin et al., 2008) and since snoRNAs molecules can be cleaved to form snoRNA fragments (snoRFs) by enzymes that remains to be elucidated (Falaleeva and Stamm, 2013; Światowy and Jagodzińśki, 2018), we studied a potential role of RNase P in snoRFs biogenesis (Sup.Fig.4).

snoRFs size distributions showed that snoRFs are highly increased in *Rpp30^18.2^* homozygous mutants, with snoRFs mostly ranging between 15 and 23nt (Sup.Fig.4A). Since there are two main snoRNA populations (box H/ACA and C/D Sup.Fig.4B), we analyzed them all together and separately to observe snoRFs coverages. Indeed, total snoRFs coverage shows that in control flies (*white*- and rescued *Rpp30^18.2^; Rpp30GFP*) there is almost no snoRFs formation (Sup.Fig.4C, upper panel, black and purple lines). However, snoRFs are highly increased in *Rpp30^18.2^* homozygous mutants (red line), mostly in 3’ of snoRNA molecules. Indeed, there is a strong accumulation of box C/D snoRFs mostly at 3’ of the snoRNA molecule, and an increase of box H/ACA 5’ and 3’ snoRFs in *Rpp30* mutant compared to controls. Sequence specificity for box C/D or H/ACA could be observed by analyzing the Logo (Sup.Fig.4D).

In contrast to *Rpp30* mutants, heterozygous and homozygous ovaries for *CG7009\** show similar profiles, with fragments mostly ranging from 21-28nt (Sup.Fig.4A). Indeed, in comparison to *white*- where almost no snoRFs could be seen, we observed that snoRFs accumulate in tRNA methylation mutant genetic backgrounds, mostly in the 3’ part of snoRNA molecules (Sup.Fig.4A and C). These results indicate that one or two alleles mutant for *CG7009\** would affect snoRNA fragmentation in a similar way. However, double mutants *CG7009*, CG5220\** have a slight increase of 5’ snoRFs and less 3’ snoRFs than *CG7009\**. In conclusion, CG7009 and CG5220 mutations lead to snoRFs accumulation, mostly derived from snoRNAs box C/D, suggesting that CG7009 and CG5220 could be important to avoid snoRFs fragmentation.

## DISCUSSION

This study describes a bioinformatic workflow to analyze tRFs populations by using Illumina-generated small RNA libraries from control and mutant flies for two key events of tRNA biology: tRNA processing and tRNA Nm methylation at the anticodon loop. We have created a new genome reference in which we included upstream and downstream sequences to mature tRNA genome sequences, and we have bioinformatically added a CCA tag, being able to analyze Type-I 3’CCA-tRFs and 5’tRFs, itRFs, Type-II, taRFs and spanners (Fig.1A).

Despite several reports published during the last years describing tRFs in different organisms, a standardized procedure to exhaustively analyze this small RNA populations, in particular using already available small RNA libraries is still lacking. Indeed, each study uses workflows with different parameters, including small RNA preparation, the number of permitted mismatches (0-3 mismatches), the length of analyzed reads, or the way the genome reference has been built (Wang et al., 2013; Goodarzi et al., 2015; Karaiskos et al., 2015a, 2015b; Kumar et al., 2015; Grigoriev and Karaiskos, 2016; Göktaş et al., 2017; Schorn et al., 2017; Shen et al., 2018; Siira et al., 2018; Kuscu et al., 2018; Liu et al., 2018; Su et al., 2019; Guan et al., 2019). Some studies have modified the genome reference to have pre-tRNAs sequences, but the number of downstream sequences can vary; some have removed introns from the genome reference; some studies have added CCA tag to the genomic reference to recover 3’CCA-tRFs but others have allowed 3 mismatches, then trimmed the CCA tag from reads to match them against the mature tRNA genome reference. Consequently, several studies are lacking some tRFs populations (3’CCA-tRFs, Type-II, spanner, taRFs). We thus designed a “user-friendly” Galaxy workflow, working with available small/mid RNA datasets, and which can be analyzed by most laboratories. We hope this workflow will provide a better understanding of tRFs biogenesis.

Using mutant flies for the RNAse P subunit (*Rpp30^18.2^*) we observed an important decrease in Type-II tRFs (Fig.9). Type-II tRFs are the consequence of RNase Z cleavage of pre-tRNAs. Interestingly, it has been described in *Drosophila* and other species that RNase P cleaves the 5’ trailer before RNase Z cleaves the 3’ trailer (Dubrovsky et al., 2004; Xie et al., 2013). In this way, an upstream defect on 5’ cleavage due to *Rpp30* mutation could affect RNase Z cleavage, thereby explaining why Type-II tRFs decrease in *Rpp30* mutants. Moreover, *Rpp30^18.2^* mutants show an accumulation of 5’tRFs. It is possible that a lack of 5’ leader cleavage affects tRNA secondary structure, promoting cleavage in the D-loop to form 5’tRFs by Dicer or other endonucleases as already shown in mammals (Li et al., 2018). Finally, 3’CCA-tRFs also increase in *Rpp30^18.2^* mutants. CCA is known to be added on mature tRNA, which suggests that Rpp30 mutation somehow affects tRNA cleavage after the CCA tRNA editing. Since 3’CCA-tRFs have been involved in TEs silencing control (Martinez et al., 2017; Schorn et al., 2017), increasing this tRFs population by promoting tRNA cleavage at the T-loop could be a way to control TEs when the main piRNA pathway is compromised. Thus, tRFs could be a versatile and adaptive source to protect genome integrity.

Besides tRFs, we observed that snoRNAs fragments (snoRFs) highly accumulate in *Rpp30* homozygous mutant ovaries. In this sense, it has been shown that snoRNAs can be a target of RNase P in some species during snoRNA maturation (Coughlin et al., 2008; Marvin et al., 2011). We know now that snoRNAs molecules can be cleaved into snoRNA fragments (snoRFs) but the enzyme(s) responsible for their cleavage remains poorly studied (Światowy and Jagodzińśki, 2018). snoRFs are aberrantly present in several pathologies such as cancer and neurodegeneration diseases (Falaleeva and Stamm, 2013; Patterson et al., 2017; Romano et al., 2017; Światowy and Jagodzińśki, 2018). Thus, RNase P could somehow participate in limiting snoRNA fragmentation to preserve homeostasis. Interestingly, mouse mutants for RNase Z (tRNA 3’ trailer processing) show an increase in snoRNA expression, believed to compensate translation defects produced by a lack of correct 3’ tRNA processing (Siira et al., 2018), but a role in snoRFs formation it has not been studied.

We also analyzed the tRFs populations in ovaries mutant for two Nm MTases of the anticodon loop of some tRNAs. Intriguingly, *CG7009\** mutants show a different tRFs profile compared to *Rpp30^18.2^* mutants (Fig.9): Type-II tRFs are decreased compared to control, Type-I 5’tRFs are slightly decreased, whereas Type-I 3’CCA tRFs are increased. On the one hand, CG7009 has been shown to methylate tRNAs at the wobble position 34 of the anticodon region and leads to an accumulation of tRNA halves fragments (around 35nt length) (M. Angelova, 2019). Thus, an accumulation of longer tRFs could impede a cleavage in the D-loop, explaining a decrease in 5’tRFs. However, in this study we cannot detect tRNA halves since our banks contain RNAs of 15-29nt. Moreover, 3’CCA tRFs increase in *CG7009\** homozygous mutants, suggesting that tRNA Nm methylation at position 34 could somehow limit T-loop cleavage. On the other hand, CG5220 has been shown to methylate position 32 of the anticodon region (M. Angelova, 2019). Interestingly, double mutants *CG7009*, CG5220\** show a different tRFs profile when compared to *CG7009\** single mutant. This result suggests that a lack of methylation in the anticodon loop region can somehow protect and/or stabilize some tRFs, as proposed recently for tRNA halves in Nm mutants (M. Angelova, 2019).

We also observed an increase of mitochondrial derived tRFs in *Rpp30^18.2^* mutants and in double mutants *CG7009*, CG5220\** when compared to control, whereas control conditions show very low levels of mito-tRFs, indicating that tRNA processing and tRNA Nm methylation pathways of the anticodon loop limit aberrant fragmentation of mitochondrial tRNAs. Mito-tRNAs are polycistronic sequences cleaved by conserved mitochondrial RNase P and Z complexes in several species (Jarrous and Gopalan, 2010; Rossmanith, 2012). Intriguingly, it has been recently shown that there is an interplay between RNase P complex and mito-tRNA methylation enzymes in human cells. Indeed, they show that mito-RNAse P recognizes, cleaves and methylates some mitochondrial tRNAs *in vitro*, which is enhanced by its interaction with a tRNA methylation cofactor (Karasik et al., 2019). Mito-RNAse P and Z dysfunction has been linked to several human mitochondrial diseases, as myopathies and neurodevelopmental disorders (Barchiesi and Vascotto, 2019; Saoura et al., 2019). Describing mitochondrial tRFs biogenesis thus help to better understand the molecular mechanisms linked to these pathologies. Interestingly, tRFs have been shown to be present in the brain of different species, and tRFs population can vary with age in *Drosophila* (Karaiskos et al., 2015a, 2015b; Grigoriev and Karaiskos, 2016; Angelova et al., 2018).

It is known that high throughput Illumina sequencing of small RNA libraries could introduce biases in tRFs detection, since tRNAs are highly modified molecules and very few techniques are able to properly describe these modifications in a transcriptome-wide way, such as ARM-seq (Cozen et al., 2015). However, a huge number of datasets are already available with valuable information to extract. By analyzing different mutants from distinct pathways we will be able to increase our knowledge on tRFs biogenesis pathway and the interactions between these different mechanisms. For example, it has been recently shown that snoRNAs association with fibrillarin can cause tRNA-CAT 2’-O-methylation at position 34 in mammalian cells, similarly to CG7009 (Vitali and Kiss, 2019). Conversely, tRNA methylation could have an impact on snoRNAs biogenesis as observed in this study. Thus, this workflow will help to analyze past, present and future small RNA sequences obtained by different means. It will be interesting in the future to obtain a tRF cartography in different tissues, organs and different species, to determine tRFs biogenesis, targets and their functions in gene expression regulation. It will be also very interesting to compare datasets obtained from classical Illumina sequencing with other techniques such as ARM-seq, which provides a read out of some modifications and may reveal additional tRFs population. Hopefully, our study will participate in the discovery of specific important nuclear tRFs or mito-tRFs that could help to understand the etiology of a wide range of human pathologies.

---

**Sup.Fig.1.**
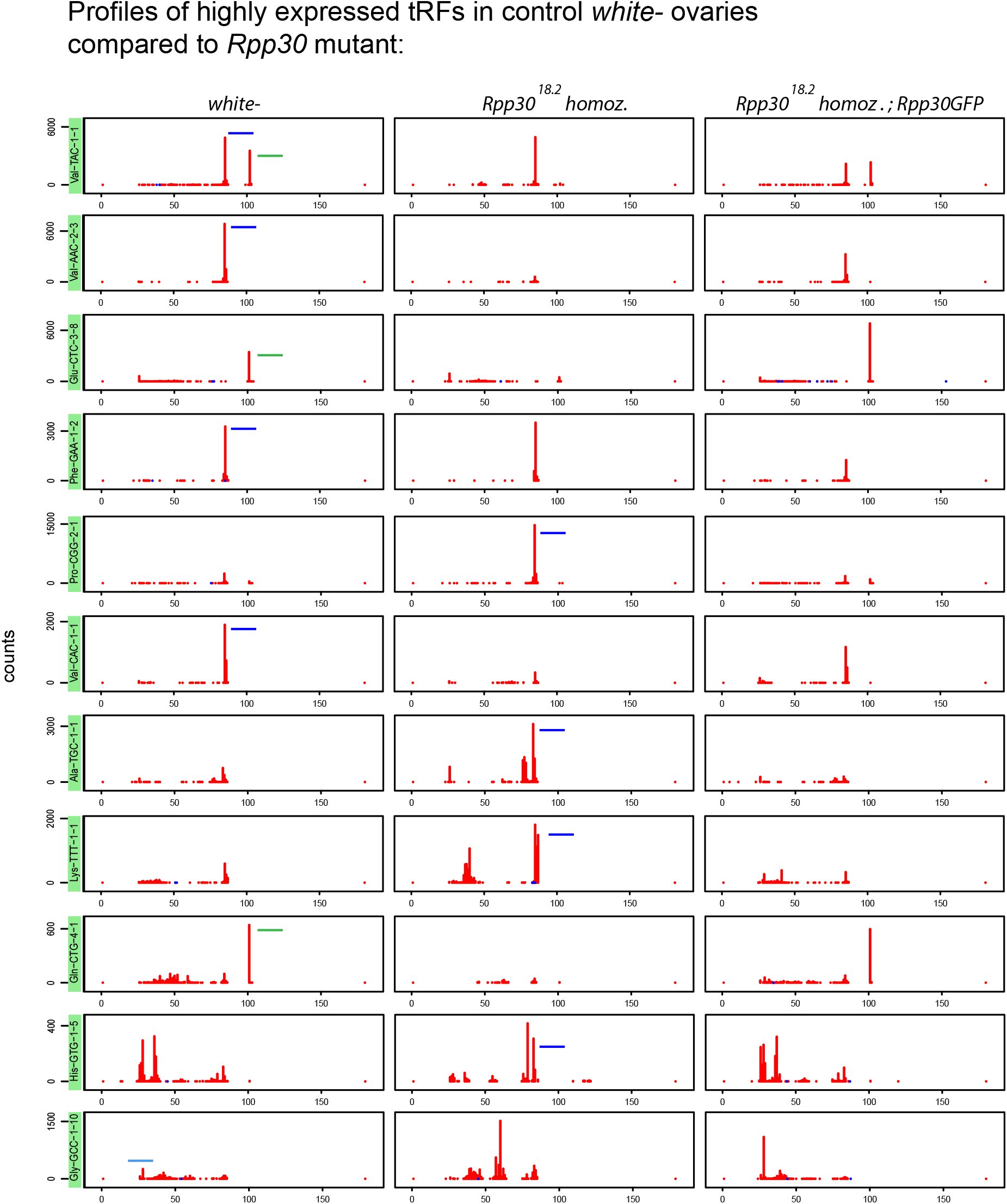
Rpp30 mutations affects tRFs biogenesis: 11 tRFs maps profiles belonging to examples of the most expressed tRFs from *white*- genotype (found in Fig.2E) are shown for the different genotypes. Maps were obtained using normalizing factors (see Methods). Since pre-tRNAs sequences are included in the tRNA genome reference, 5’tRFs are located at the position 25nt instead of position 0nt. 3’CCA-tRFs are located around the position 75nt and Type-II tRFs are located around position 100nt, depending on the length of the tRNA and if they have an intron. The peak determines the beginning of the sequence. tRFs categories are schematized in *white*- and *Rpp30^18.2^* homozygous for better comparison: 5’tRFs in light blue, 3’CCA in dark blue and Type-II in green.

**Sup.Fig.2.**
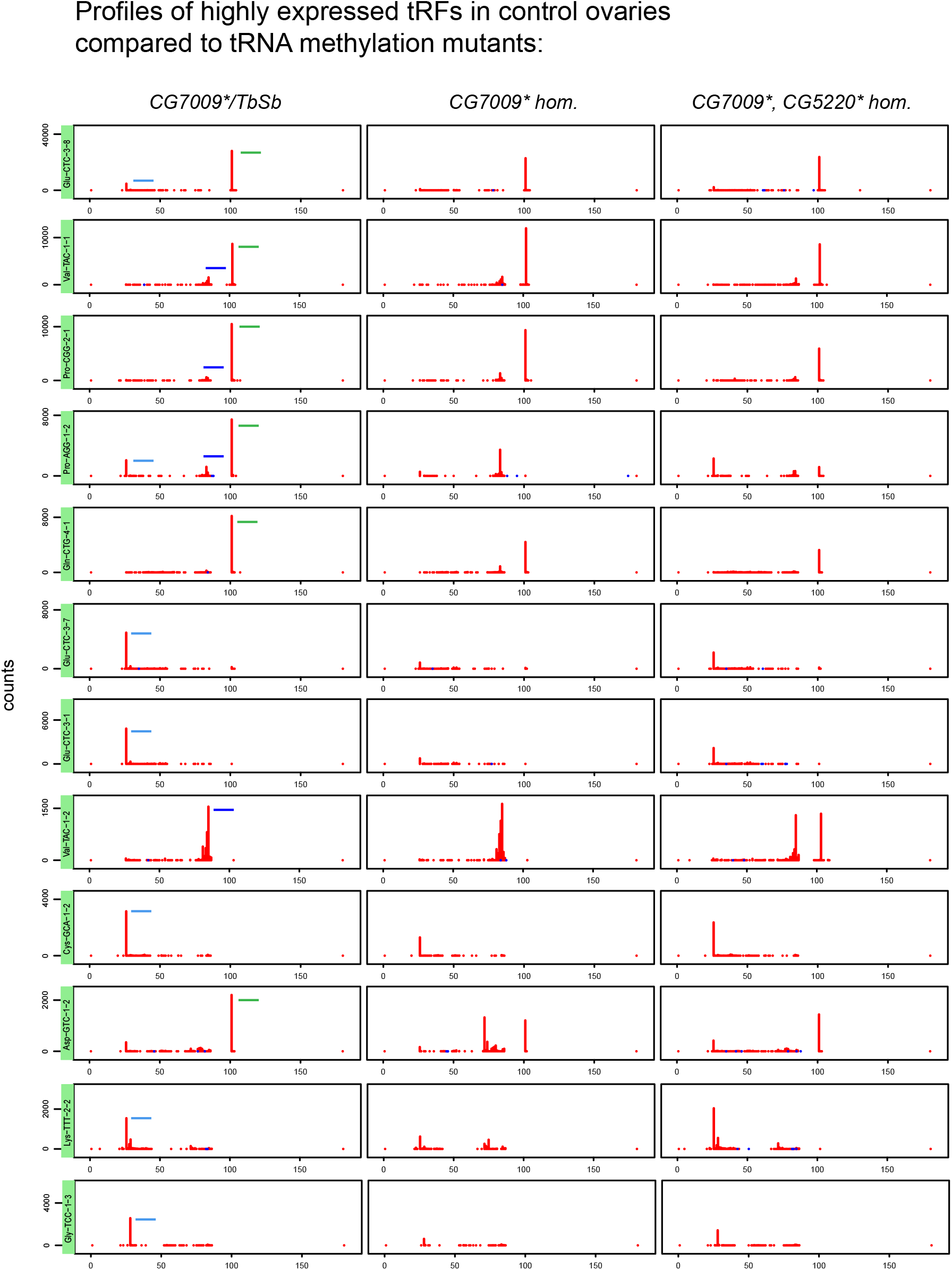
tRNA methylation defects alters tRFs biogenesis: 12 tRFs normalized maps profiles belonging to examples of the most changed (decreased or increased) tRFs from *CG7009\** homozygous/heterozygous ratio (see Fig.7B) are shown for the different genotypes. Since pre-tRNAs sequences are included in the tRNA-reference, 5’tRFs are located at the position 25nt instead of position 0nt. 3’CCA-tRFs are located around the position 75nt and Type-II tRFs are located around position 100nt, depending on the length of the tRNA and if they have an intron*. The peak determines the beginning of the sequence. tRFs categories are schematized in *CG7009*/Tb,Sb* heterozygous and homozygous mutants for better comparison: 5’tRFs in light blue, 3’CCA in dark blue and Type-II in green. Ratio’s values above 1 : tRFs increased in *CG7009\** homozygous mutants. Ratio’s values below 1: tRFs decreased in *CG7009\** homozygous mutants.

**Sup.Fig.3.**
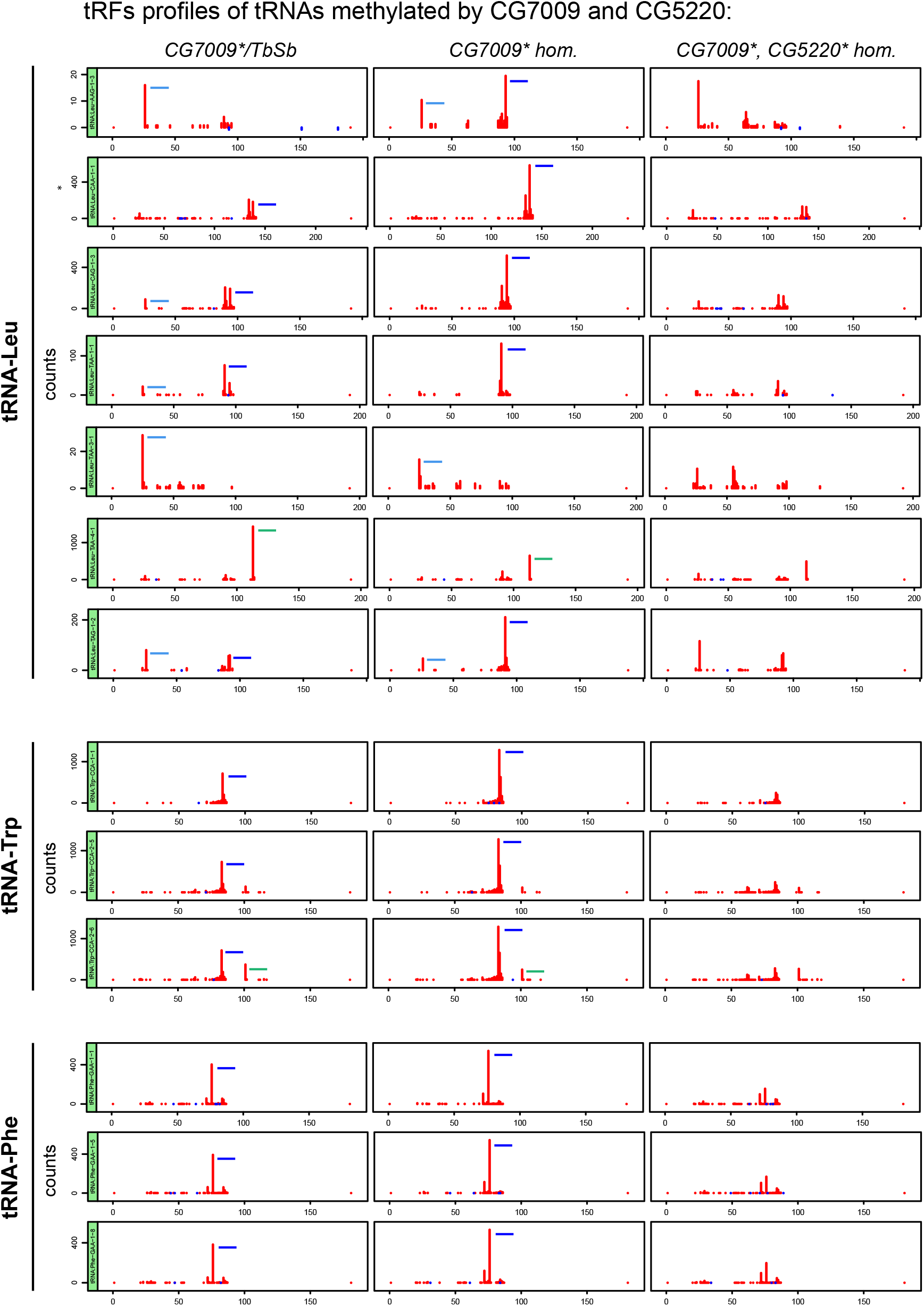
tRNA methylation defects alters tRFs biogenesis: 13 tRFs normalized profiles belonging to examples of tRNA targets of CG7009 and CG5220 are shown for the different genotypes. Since pre-tRNAs sequences are included in the tRNA-reference, 5’tRFs are located at the position 25nt instead of position 0nt. 3’CCA-tRFs are located around the position 75nt and Type-II tRFs are located around position 100nt, depending on the length of the tRNA and if they have an intron. The peak determines the beginning of the sequence tRFs are schematized in *CG7009*/Tb,Sb* heterozygous and homozygous mutants for better comparison: 5’tRFs in light blue, 3’CCA in dark blue and Type-II in green.

**Sup.Fig.4.**
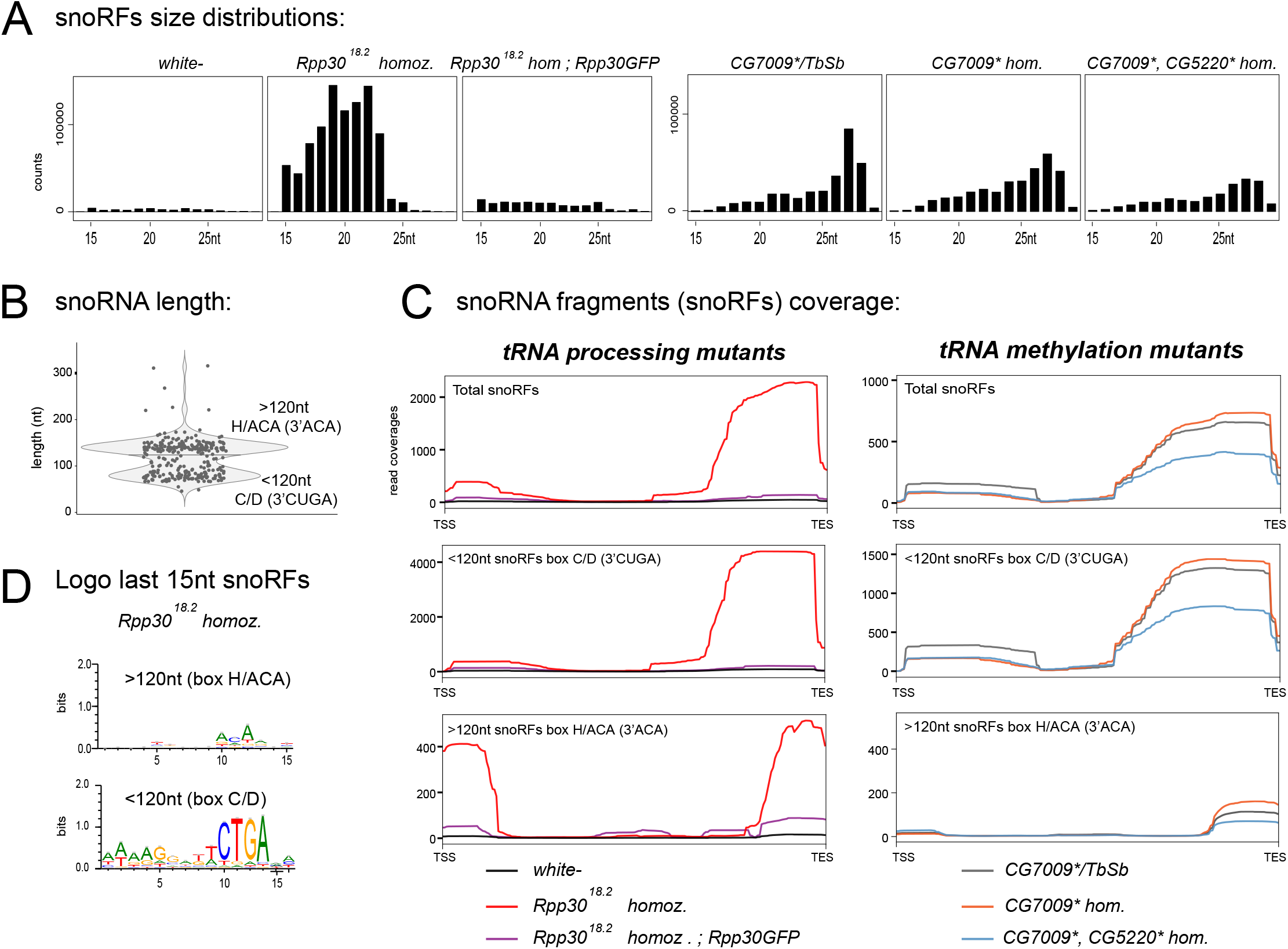
tRNA methylation affects snoRNA fragments (snoRFs). **A.** General size distribution (15-29nt) of normalized snoRFs read counts is shown for the different genotypes of tRNA processing and tRNA methylation mutants. **B.** A violin plot reflects snoRNAs populations found in *Drosophila melanogaster* genome. snoRNAs of more than 120nt belong to *box H/ACA* class whereas snoRNAs of less than 120nt belong to *box C/D* class. **C.** snoRFs coverage (scaling factors used, see Methods) is shown. TSS: Transcription Start Site. TES, Transcription End Site. **D.** Logo for the most representative sequences found in the last 15nt of snoRFs is shown for *Rpp30* mutant (issued from *fasta* files containing all different sequences).

**Sup.Fig.5.**
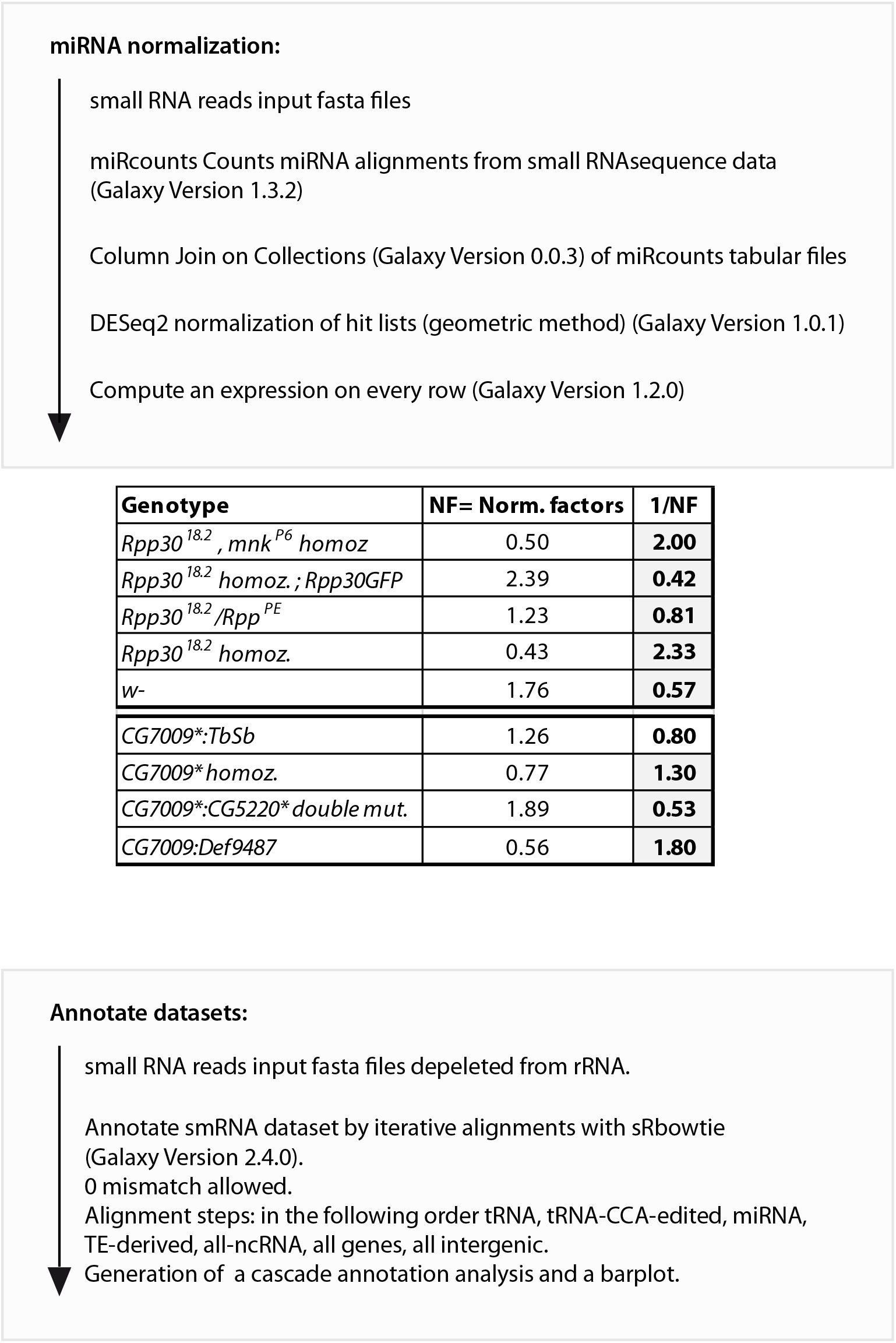
Workflow related to miRNA normalization, scale factors and and global cascade annotation. The Workflow to obtain Scale Factors is detailed. Scale factors for all the genotypes used in this study are shown (Molla-Herman et al., 2015; M. Angelova, 2019). For simplicity, only *white-, Rpp30^18.2^, Rpp30^18.2^; ubiRpp30GFP, CG7009\** heterozygous, *CG7009\** homozygous and *CG7009*, CG5220\** double mutants have been used for the main figures. The Workflow related to Fig.3 and 6 (cascade annotation) is detailed.

**Sup.Fig.6.**
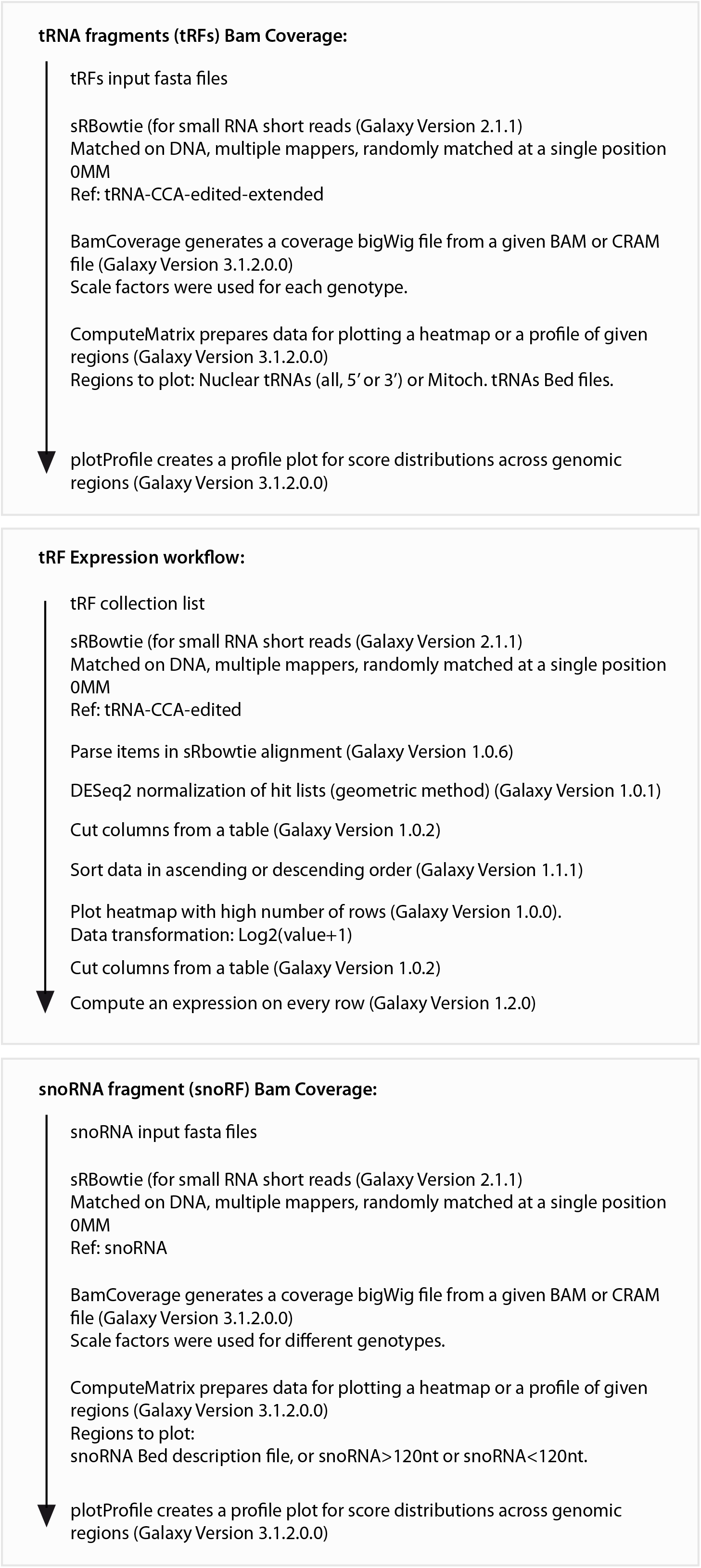
Workflow related to Bam Coverages and tRFs expression heatmaps. Workflows related to Fig. 3 and 6 (tRFs Bam Coverages), related to Fig.2, 4 and 7 (tRFs Heatmap) and related to Sup.Fig.4 (snoRFs Bam Coverage) are detailed.

## Acknowledgements

We thank Ana Maria Vallés (Collège de France, Paris) for helping with manuscript reading.

## Author contributions

Anahí Mollá Herman (AMH). Margarita T. Angelova (MTA). Clément Carré (CC). Christophe Antoniewski (CA). Jean-René HUYNH (JRH): AMH and MTA performed experiments. AMH and CA designed and performed bioinformatic data analyses. AMH wrote the manuscript. JRH participated in data analysis and manuscript writing. MTA, CA and CC participated in manuscript writing.

JRH lab is supported by CNRS, Inserm, Collège de France, FRM (Equipe FRM DEQ20160334884), ANR (ANR-13-BSV2-0007-02 PlasTiSiPi; ANR-15-CE13-0001-01, AbsCyStem) and Bettencourt-Schueller foundations. CC received financial support from CNRS and Sorbonne Université, Fondation Maladies Rares (FMR), IBPS “AI 2018” and the COST action EPITRAN CA16120. CC is a member of the COST action “EPITRAN” CA16120.

## Bibliography

Angelova, M. T., Dimitrova, D. G., Dinges, N., Lence, T., Worpenberg, L., Carré, C., et al. (2018). The emerging field of epitranscriptomics in neurodevelopmental and neuronal disorders. Front. Bioeng. Biotechnol. 6, 1–15. doi:10.3389/fbioe.2018.00046.

Angrisani, A., Tafer, H., Stadler, P. F., and Furia, M. (2015). Developmentally regulated expression and expression strategies of Drosophila snoRNAs. Insect Biochem. Mol. Biol. 61, 69–78. doi:10.1016/j.ibmb.2015.01.013.

Balatti, V., Nigita, G., Veneziano, D., Drusco, A., Stein, G. S., Messier, T. L., et al. (2017). tsRNA signatures in cancer. Proc. Natl. Acad. Sci. U. S. A. 114, 8071–8076. doi:10.1073/pnas.1706908114.

Barchiesi, A., and Vascotto, C. (2019). Transcription, processing, and decay of mitochondrial RNA in health and disease. Int. J. Mol. Sci. 20. doi:10.3390/ijms20092221.

Coughlin, D. J., Pleiss, J. A., Walker, S. C., Whitworth, G. B., and Engelke, D. R. (2008). Genome-wide search for yeast RNase P substrates reveals role in maturation of intron-encoded box C/D small nucleolar RNAs. Proc. Natl. Acad. Sci. U. S. A. 105, 12218–12223. doi:10.1073/pnas.0801906105.

Cozen, A. E., Quartley, E., Holmes, A. D., Hrabeta-Robinson, E., Phizicky, E. M., and Lowe, T. M. (2015). ARM-seq: AlkB-facilitated RNA methylation sequencing reveals a complex landscape of modified tRNA fragments. Nat. Methods. doi:10.1038/nmeth.3508.

Czech, B., Munafò, M., Ciabrelli, F., Eastwood, E. L., Fabry, M. H., Kneuss, E., et al. (2018). piRNA-Guided Genome Defense: From Biogenesis to Silencing. Annu. Rev. Genet. doi:10.1146/annurev-genet-120417-031441.

Dai, Q., Zheng, G., Schwartz, M. H., Clark, W. C., and Pan, T. (2017). Selective Enzymatic Demethylation of N2,N2-Dimethylguanosine in RNA and Its Application in High-Throughput tRNA Sequencing. Angew. Chemie - Int. Ed. 56, 5017–5020. doi:10.1002/anie.201700537.

Delaunay, S., and Frye, M. (2019). RNA modifications regulating cell fate in cancer. Nat. Cell Biol. 21, 552–559. doi:10.1038/s41556-019-0319-0.

Dimitrova, D. G., Teysset, L., and Carré, C. (2019). RNA 2’-O-Methylation (Nm) modification in human diseases. Genes (Basel). 10, 1–23. doi:10.3390/genes10020117.

Dubrovsky, E. B., Dubrovskaya, V. A., Levinger, L., Schiffer, S., and Marchfelder, A. (2004). Drosophila Rnase Z processes mitochondrial and nuclear pre-tRNA 3’ ends in vivo. Nucleic Acids Res. 32, 255–262. doi:10.1093/nar/gkh182.

Falaleeva, M., and Stamm, S. (2013). Processing of snoRNAs as a new source of regulatory non-coding RNAs: SnoRNA fragments form a new class of functional RNAs. BioEssays 35, 46–54. doi:10.1002/bies.201200117.

Falaleeva, M., Welden, J. R., Duncan, M. J., and Stamm, S. (2017). C/D-box snoRNAs form methylating and non-methylating ribonucleoprotein complexes: Old dogs show new tricks. BioEssays 39, 1–15. doi:10.1002/bies.201600264.

Genenncher, B., Durdevic, Z., Hanna, K., Zinkl, D., Mobin, M. B., Senturk, N., et al. (2018). Mutations in Cytosine-5 tRNA Methyltransferases Impact Mobile Element Expression and Genome Stability at Specific DNA Repeats. Cell Rep. 22, 1861–1874. doi:10.1016/j.celrep.2018.01.061.

Göktaş, Ç., Yiğit, H., Coşacak, M. İ., and Akgül, B. (2017). Differentially expressed tRNA-derived small RNAs co-sediment primarily with non-polysomal fractions in Drosophila. Genes (Basel). 8. doi:10.3390/genes8110333.

Goodarzi, H., Liu, X., Nguyen, H. C. B., Zhang, S., Fish, L., and Tavazoie, S. F. (2015). Endogenous tRNA-derived fragments suppress breast cancer progression via YBX1 displacement. Cell 161, 790–802. doi:10.1016/j.cell.2015.02.053.

Grigoriev, A., and Karaiskos, S. (2016). Dynamics of tRNA fragments and their targets in aging mammalian brain. F1000Research 5, 1–16. doi:10.12688/f1000research.10116.1.

Guan, L., Karaiskos, S., and Grigoriev, A. (2019). Inferring targeting modes of Argonaute-loaded tRNA fragments. RNA Biol. 00, 1–11. doi:10.1080/15476286.2019.1676633.

Guy, M. P., Shaw, M., Weiner, C. L., Hobson, L., Stark, Z., Rose, K., et al. (2015). Defects in tRNA Anticodon Loop 2’-O-Methylation Are Implicated in Nonsyndromic X-Linked Intellectual Disability due to Mutations in FTSJ1. Hum. Mutat. doi:10.1002/humu.22897.

Haeusler, R. A., and Engelke, D. R. (2006). Survey and Summary: Spatial organization of transcription by RNA polymerase III. Nucleic Acids Res. doi:10.1093/nar/gkl656.

Huang, Z. P., Zhou, H., Hua-Liang, H., Chen, C. L., Liang, D., and Liang-Hu, Q. (2005). Genome-wide analyses of two families of snoRNA genes from Drosophila melanogaster, demonstrating the extensive utilization of introns for coding of snoRNAs. Rna 11, 1303–1316. doi:10.1261/rna.2380905.

Jarrous, N. (2017). Roles of RNase P and Its Subunits. Trends Genet. 33, 594–603. doi:10.1016/j.tig.2017.06.006.

Jarrous, N., and Gopalan, V. (2010). Archaeal/Eukaryal RNase P: Subunits, functions and RNA diversification. Nucleic Acids Res. 38, 7885–7894. doi:10.1093/nar/gkq701.

Jordan Ontiveros, R., Stoute, J., and Liu, K. F. (2019). The chemical diversity of RNA modifications. Biochem. J. 476, 1227–1245. doi:10.1042/BCJ20180445.

Karaiskos, S., Naqvi, A. S., Swanson, K. E., and Grigoriev, A. (2015a). Age-driven modulation of tRNA-derived fragments in Drosophila and their potential targets. Biol. Direct 10, 1–14. doi:10.1186/s13062-015-0081-6.

Karaiskos, S., Naqvi, A. S., Swanson, K. E., and Grigoriev, A. (2015b). Age-driven modulation of tRNA-derived fragments in Drosophila and their potential targets. Biol. Direct 10, 1–13. doi:10.1186/s13062-015-0081-6.

Karasik, A., Fierke, C. A., and Koutmos, M. (2019). Interplay between substrate recognition, 5’ end tRNA processing and methylation activity of human mitochondrial RNase P. RNA. doi:10.1261/rna.069310.118.

Kumar, P., Kuscu, C., and Dutta, A. (2016). Biogenesis and Function of Transfer RNA-Related Fragments (tRFs). Trends Biochem. Sci. 41, 679–689. doi:10.1016/j.tibs.2016.05.004.

Kumar, P., Mudunuri, S. B., Anaya, J., and Dutta, A. (2015). tRFdb: A database for transfer RNA fragments. Nucleic Acids Res. 43, D141–D145. doi:10.1093/nar/gku1138.

Kuscu, C., Kumar, P., Kiran, M., Su, Z., Malik, A., and Dutta, A. (2018). tRNA fragments (tRFs) guide Ago to regulate gene expression post-transcriptionally in a Dicer-independent manner. Rna 24, 1093–1105. doi:10.1261/rna.066126.118.

Li, L., Gu, W., Liang, C., Liu, Q., Mello, C. C., and Liu, Y. (2012). The translin-TRAX complex (C3PO) is a ribonuclease in tRNA processing. Nat. Struct. Mol. Biol. 19, 824–830. doi:10.1038/nsmb.2337.

Li, S., Xu, Z., and Sheng, J. (2018). tRNA-derived small RNA: A novel regulatory small non-coding RNA. Genes (Basel). 9. doi:10.3390/genes9050246.

Liu, S., Chen, Y., Ren, Y., Zhou, J., Ren, J., Lee, I., et al. (2018). A tRNA-derived RNA Fragment Plays an Important Role in the Mechanism of Arsenite -induced Cellular Responses. Sci. Rep. 8, 1–9. doi:10.1038/s41598-018-34899-2.

M. Angelova, et al. (2019). tRNA 2’-O-methylation modulates small RNA silencing and life span in. bioRxiv. Available at: https://www.biorxiv.org/content/10.1101/699934v2.

Martinez, G., Choudury, S. G., and Slotkin, R. K. (2017). TRNA-derived small RNAs target transposable element transcripts. Nucleic Acids Res. 45, 5142–5152. doi:10.1093/nar/gkx103.

Marvin, M. C., Clauder-Münster, S., Walker, S. C., Sarkeshik, A., Yates, J. R., Steinmetz, L. M., et al. (2011). Accumulation of noncoding RNA due to an RNase P defect in Saccharomyces cerevisiae. Rna 17, 1441–1450. doi:10.1261/rna.2737511.

Molla-Herman, A., Vallés, A. M., Ganem-Elbaz, C., Antoniewski, C., and Huynh, J. (2015). tRNA processing defects induce replication stress and Chk2-dependent disruption of piRNA transcription. EMBO J. 34, 3009–3027. doi:10.15252/embj.201591006.

Motorin, Y., and Helm, M. (2019). Methods for RNA modification mapping using deep sequencing: Established and new emerging technologies. Genes (Basel). doi:10.3390/genes10010035.

Park, O. H., Ha, H., Lee, Y., Boo, S. H., Kwon, D. H., Song, H. K., et al. (2019). Endoribonucleolytic Cleavage of m6A-Containing RNAs by RNase P/MRP Complex. Mol. Cell 74, 494–507.e8. doi:10.1016/j.molcel.2019.02.034.

Patterson, D. G., Roberts, J. T., King, V. M., Houserova, D., Barnhill, E. C., Crucello, A., et al. (2017). Human snoRNA-93 is processed into a microRNA-like RNA that promotes breast cancer cell invasion. npj Breast Cancer 3, 1–12. doi:10.1038/s41523-017-0032-8.

Pintard, L., Lecointe, F., Bujnicki, J. M., Bonnerot, C., Grosjean, H., and Lapeyre, B. (2002). Trm7p catalyses the formation of two 2’-O-methylriboses in yeast tRNA anticodon loop. EMBO J. doi:10.1093/emboj/21.7.1811.

Romano, G., Veneziano, D., Acunzo, M., and Croce, C. M. (2017). Small non-coding RNA and cancer. Carcinogenesis 38, 485–491. doi:10.1093/carcin/bgx026.

Rossmanith, W. (2012). Of P and Z: Mitochondrial tRNA processing enzymes. Biochim. Biophys. Acta - Gene Regul. Mech. 1819, 1017–1026. doi:10.1016/j.bbagrm.2011.11.003.

Saoura, M., Powell, C. A., Kopajtich, R., Alahmad, A., AL-Balool, H. H., Albash, B., et al. (2019). Mutations in ELAC2 associated with hypertrophic cardiomyopathy impair mitochondrial tRNA 3’-end processing. Hum. Mutat. 40, 1731–1748. doi:10.1002/humu.23777.

Schaefer, M., Pollex, T., Hanna, K., Tuorto, F., Meusburger, M., Helm, M., et al. (2014). Course Numbering Grades Grade Notations. 1590–1595. doi:10.1101/gad.586710.1590.

Schorn, A. J., Gutbrod, M. J., LeBlanc, C., and Martienssen, R. (2017). LTR-Retrotransposon Control by tRNA-Derived Small RNAs. Cell 170, 61–71.e11. doi:10.1016/j.cell.2017.06.013.

Schorn, A. J., and Martienssen, R. (2018). Tie-Break: Host and Retrotransposons Play tRNA. Trends Cell Biol. 28, 793–806. doi:10.1016/j.tcb.2018.05.006.

Shen, Y., Yu, X., Zhu, L., Li, T., Yan, Z., and Guo, J. (2018). Transfer RNA-derived fragments and tRNA halves: biogenesis, biological functions and their roles in diseases. J. Mol. Med. 96, 1167–1176. doi:10.1007/s00109-018-1693-y.

Siira, S. J., Rossetti, G., Richman, T. R., Perks, K., Ermer, J. A., Kuznetsova, I., et al. (2018). Concerted regulation of mitochondrial and nuclear non-coding RNA s by a dual-targeted RN ase Z. EMBO Rep. 19, 1–18. doi:10.15252/embr.201846198.

Sokołowski, M., Klassen, R., Bruch, A., Schaffrath, R., and Glatt, S. (2018). Cooperativity between different tRNA modifications and their modification pathways. Biochim. Biophys. Acta - Gene Regul. Mech. 1861, 409–418. doi:10.1016/j.bbagrm.2017.12.003.

Su, Z., Kuscu, C., Malik, A., Shibata, E., and Dutta, A. (2019). Angiogenin generates specific stress-induced tRNA halves and is not involved in tRF-3-mediated gene silencing. J. Biol. Chem. 294, 16930–16941. doi:10.1074/jbc.RA119.009272.

Sun, C., Fu, Z., Wang, S., Li, J., Li, Y., Zhang, Y., et al. (2018). Roles of tRNA-derived fragments in human cancers. Cancer Lett. 414, 16–25. doi:10.1016/j.canlet.2017.10.031.

Światowy, W., and Jagodzińśki, P. P. (2018). Molecules derived from tRNA and snoRNA: Entering the degradome pool. Biomed. Pharmacother. 108, 36–42. doi:10.1016/j.biopha.2018.09.017.

Vitali, P., and Kiss, T. (2019). Cooperative 2’-o-methylation of the wobble cytidine of human elongator tRNAmet(cat) by a nucleolar and a cajal bodyspecific box C/D RNP. Genes Dev. 33, 741–746. doi:10.1101/gad.326363.119.

Wang, Q., Lee, I., Ren, J., Ajay, S. S., Lee, Y. S., and Bao, X. (2013). Identification and functional characterization of tRNA-derived RNA fragments (tRFs) in respiratory syncytial virus infection. Mol. Ther. 21, 368–379. doi:10.1038/mt.2012.237.

Wellner, K., Betat, H., and Mörl, M. (2018). A tRNA’s fate is decided at its 3’ end: Collaborative actions of CCA-adding enzyme and RNases involved in tRNA processing and degradation. Biochim. Biophys. Acta - Gene Regul. Mech. 1861, 433–441. doi:10.1016/j.bbagrm.2018.01.012.

Willis, I. M., and Moir, R. D. (2018). Signaling to and from the RNA Polymerase III Transcription and Processing Machinery. Annu. Rev. Biochem. 87, 75–100. doi:10.1146/annurev-biochem-062917-012624.

Xie, X., Dubrovskaya, V., Yacoub, N., Walska, J., Gleason, T., Reid, K., et al. (2013). Developmental roles of Drosophila tRNA processing endonuclease RNase ZL as revealed with a conditional rescue system. Dev. Biol. 381, 324–340. doi:10.1016/j.ydbio.2013.07.005.

Yamanaka, S., and Siomi, H. (2015). Misprocessed tRNA response targets pi RNA clusters. EMBO J. 34, 2988–2989. doi:10.15252/embj.201593322.

Zhu, L., Liu, X., Pu, W., and Peng, Y. (2018). tRNA-derived small non-coding RNAs in human disease. Cancer Lett. 419, 1–7. doi:10.1016/j.canlet.2018.01.015.

